# Data-driven prediction of colonization outcomes for complex microbial communities

**DOI:** 10.1101/2023.04.19.537502

**Authors:** Lu Wu, Xu-Wen Wang, Zining Tao, Tong Wang, Wenlong Zuo, Yu Zeng, Yang-Yu Liu, Lei Dai

## Abstract

Complex microbial interactions can lead to different colonization outcomes of exogenous species, be they pathogenic or beneficial in nature. Predicting the colonization of exogenous species in complex communities remains a fundamental challenge in microbial ecology, mainly due to our limited knowledge of the diverse physical, biochemical, and ecological processes governing microbial dynamics. Here, we proposed a data-driven approach independent of any dynamics model to predict colonization outcomes of exogenous species from the baseline compositions of microbial communities. We systematically validated this approach using synthetic data, finding that machine learning models (including Random Forest and neural ODE) can predict not only the binary colonization outcome but also the post-invasion steady-state abundance of the invading species. Then we conducted colonization experiments for two commensal gut bacteria species *Enterococcus faecium* and *Akkermansia muciniphila* in hundreds of human stool-derived *in vitro* microbial communities, confirming that the data-driven approach can successfully predict the colonization outcomes. Furthermore, we found that while most resident species were predicted to have a weak negative impact on the colonization of exogenous species, strongly interacting species could significantly alter the colonization outcomes, e.g., the presence of *Enterococcus faecalis* inhibits the invasion of *E. faecium*. The presented results suggest that the data-driven approach is a powerful tool to inform the ecology and management of complex microbial communities.

## Introduction

Microbial communities are constantly exposed to the invasion of exogenous species, which can significantly alter their composition and function (*1-4*). The capacity of a microbial community to resist invasion is regarded as an emergent property (i.e., from individual parts to a holistic function) arising from the complex interactions among its constituent species (*5*). Theoretical studies have found that communities with higher diversity or stronger interaction strengths among species are more resistant to potential invaders (*6-8*), attributed to the fact that communities with higher diversity can occupy more niches and provide functional redundancy, making it more difficult for an invading species to establish and thrive.

The role of host-associated microbiota in defending against pathogens has been extensively studied (*9-12*), particularly in the context of the human gut microbiome. For the human gut microbiome, invasion of resident microbial communities can occur when non-resident bacteria from foods and the upper gastrointestinal tract reach the gut ecosystem (*13, 14*). The resident microbes in the gut ecosystem outcompete and exclude invaders through a combination of mechanisms, such as producing antimicrobial compounds (*15, 16*), competing for nutrients and space (*17-20*), and modulating the host’s immune response (*21, 22*). However, the composition of the human gut microbiota can vary significantly across individuals (*23, 24*) and over time (*25, 26*). Dietary shifts, medication, and other environmental factors can greatly alter the composition of the gut microbial community (*27, 28*). These interpersonal and dynamic variations in the gut microbiome can lead to significantly different colonization outcomes, such as resistance to pathogens (*29, 30*) and probiotics (*31, 32*). For example, antibiotics treatment often leads to a loss of diversity and promotes the invasion of exogenous species (*30, 33*). The ability to predict and alter the colonization outcomes (i.e., prevent the engraftment of pathogens and promote the engraftment of probiotics) is critical for personalized microbiota-based interventions in nutrition and medicine.

Despite the accumulating empirical studies, predicting the colonization outcomes in complex communities, such as the human gut microbiome, remains a fundamental challenge due to limited knowledge of interspecies interactions. For a meta-community of *N* species (*N* ranges from hundreds to thousands for the human gut microbiome), we would need to isolate and culture all these species (which is already a formidable task) and conduct a considerable number of experiments to map pairwise interactions (on the order of ∼*O(N*^*2*^)), not to mention higher-order interactions. Thus, novel approaches are needed to study the ecology of highly complex communities.

The key question is: can we achieve system-level predictions for complex ecological systems without requiring detailed mechanistic information? Colonization outcome can be viewed as a mapping from the community structure of a complex ecological system (i.e., the pre-invasion community profile) to its function (i.e., the post-invasion abundance of the invading species). Recently, the application of data-driven models (machine learning and deep learning) has shown great promise in predicting the emergent properties of complex biomolecules, such as protein structure (mapping from protein sequence to structure) (*34*), promoter strength (mapping from DNA sequence to function) (*35*). In contrast, this paradigm shift to data-driven approaches has received much less attention in ecology (*36*).

Here, we proposed a data-driven approach to predict colonization outcomes of exogenous species in complex microbial communities. First, we systematically evaluated the approach using synthetic data generated by classical ecological dynamical models. We found that, with sufficient sample size in training data (on the order of ∼*O(N)*), the colonization outcomes (i.e., whether an exogenous species can engraft and what its abundance would be if it can engraft) can be accurately predicted by machine learning models. Then, we generated large-scale datasets with *in-vitro* experimental outcomes of two representative species (*E. faecium* and *A. muciniphila*) colonizing human stool-derived microbial communities. We validated that machine-learning models, including Random Forest and neural ODE, can accurately predict the colonization outcomes in experiments. Finally, we used the machine learning models to infer species with large colonization impacts and experimentally demonstrated that the introduction of strongly interacting species can significantly alter the colonization outcomes. Our results suggest that the function of complex microbial communities is predictable and tunable via the data-driven approach.

## Results

### The data-driven approach of predicting colonization outcomes for complex microbial communities

Let’s consider a meta-community with a pool of *N* microbial species, denoted as Ω = {1, ⋯, *N*}. Consider a large set of *M* microbiome samples, denoted as 𝒮 = {1, …, *M*}, collected from this meta-community. A microbiome sample *s* ∈ 𝒮 can be considered as a local community of the meta-community with a subset of coexisting species (**Fig.1A**). For a local community *s*, if an exogenous species-*i* (still in the species pool Ω, but not in *s*) is introduced to *s*, whether it can successfully colonize the community or not, as well as its post-invasion abundance *x*_*i*_, will depend on the baseline composition of *s*. For example, it is easier (or harder) for species-*i* to colonize *s* if some resident species strongly promote (or inhibit) its growth of species-*i*, respectively. Hereafter, we call the community *s permissive* (or *resistant*) to species-*i* if species-*i* can (or cannot) successfully colonize *s*, respectively. If we only have the information about species-*i* and a community *s*, it may seem impossible to accurately predict the colonization outcome without detailed knowledge about microbial interactions. However, if we have access to the data from colonization experiments of many local communities, then, in principle, we can formalize the colonization outcome prediction problem as a machine-learning task that can be solved in a data-driven fashion. To ensure the problem is mathematically well-defined, we must assume that the different local communities in this meta-community share identical assembly rules and microbial interactions (*37*). This way, the colonization outcomes of some local communities can be used to train a machine-learning model to predict the colonization outcomes of other local communities.

**Figure 1.**
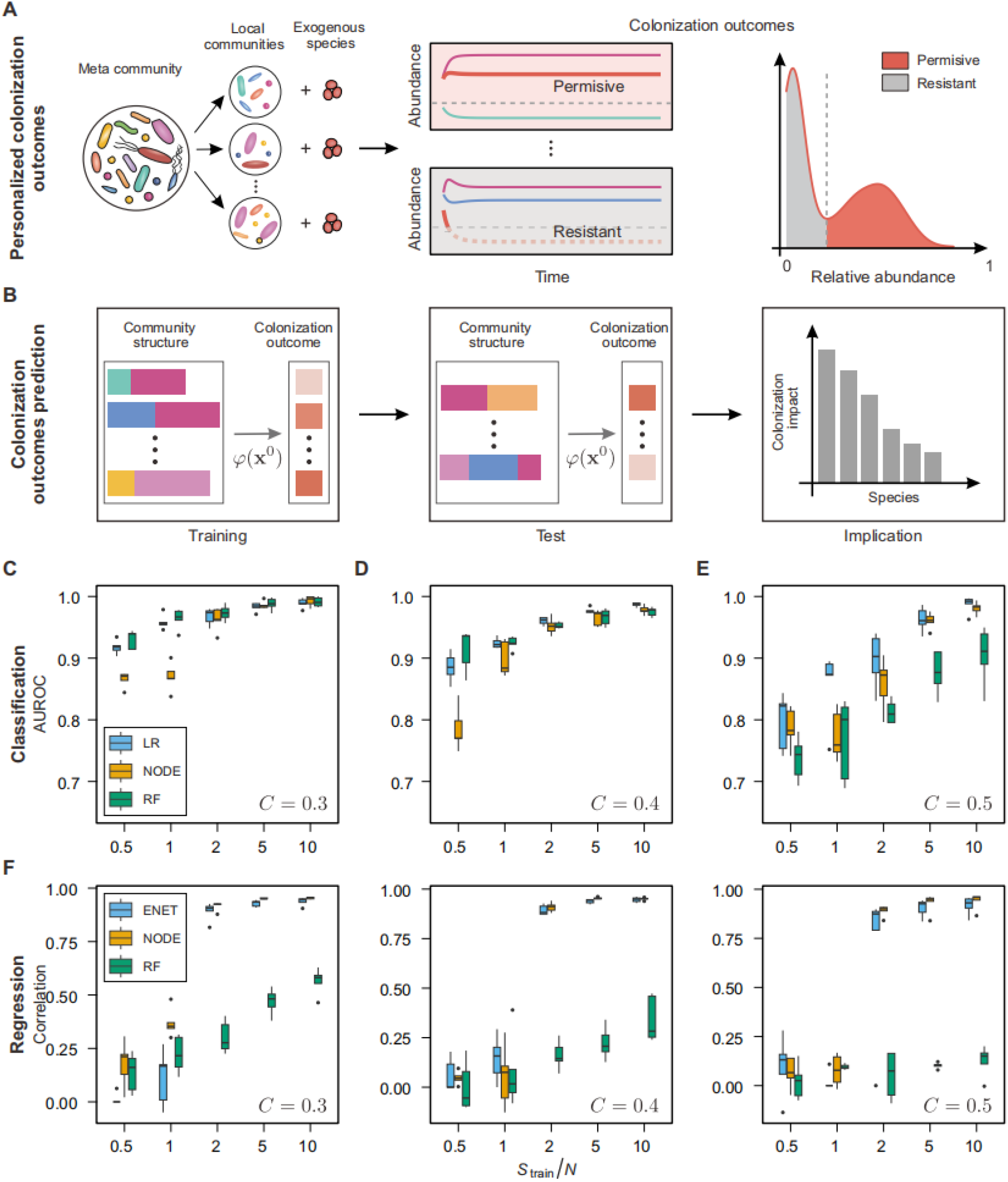
Prediction of colonization outcomes for complex microbial communities via the data-driven approach. **(A)** Each individual’s microbiome can be viewed as a local community, a subset of the meta-community of microbial species. For an exogenous species that invades the local communities, its colonization outcome (e.g., permissive or resistant) can be highly personalized, depending on the composition of local communities. **(B)** Colonization outcome prediction (COP) can be solved by learning the mapping from the baseline taxonomic profile to the post-invasion abundance of the exogenous species (i.e., *φ*: **x**^0^ ↦ x^1^). **(C-E)** Evaluation of the data-driven approach in solving the classification task of COP. AUROC of three machine learning models, including Logistic Regression (LR), COP-Neural Ordinary Differential Equations classifier (NODE), and Random Forest classifier (RF). **(F-H)** Evaluation of the data-driven approach in solving the regression task of COP. Pearson correlation between the true abundance and the abundance predicted by three machine learning models, including Elastic Net Linear Regression (ENET), COP-NODE regressor (NODE), and Random Forest regressor (RF).

Consider species-*i* as the exogenous species to a local community *s*. Note that the baseline abundance of species-*i* is zero (i.e., 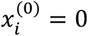) in *s* before the invasion. With some initial abundance, the exogenous species will interact with the resident species in *s*, and its post-invasion steady-state abundance is denoted as 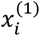. We propose to solve the Colonization Outcome Prediction (COP) problem using machine-learning models that treat the baseline (i.e., pre-invasion) taxonomic profile **x**^(0)^ as inputs and the steady state abundance of the invasive species 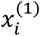 as output (**Fig.1B**). Mathematically, we intend to learn the mapping from the baseline taxonomic profile of a community **x**^(0)^ to the steady state abundance of the invading species 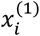, i.e., 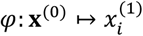. In addition, this mapping could help us infer the impact of each resident species on the colonization of the exogenous species.

We conducted *in silico* simulations to validate the feasibility of our approach. We generated synthetic data of colonization outcomes using the Generalized Lotka-Volterra (GLV) model with *N* = 100 species in the meta-community (see **Methods**). The initial species collection of each sample (i.e., a local community) consists of 30 species randomly drawn from the (*N* − 1) species pool (the exogenous species is absent in all the local communities). We generated the baseline profiles of local communities by running the GLV dynamics to a steady state. The exogenous species was then added to each local community, and its post-invasion abundance was obtained by running the GLV dynamics to a new steady state.

We can formalize COP as two sub-problems: (1) *Classification*: predict whether an exogenous species can colonize a local community; (2) *Regression*: predict the steady-state abundance of an exogenous species after colonization. Using the synthetic data generated by the GLV model, we first addressed the *classification* problem, i.e., predicting whether the invading species can colonize a community. We employed three models covering representative categories of machine learning: Logistic Regression, Random Forest classifier, and COP-Neural Ordinary Differential Equations (COP-NODE) classifier (see **Methods**). We tuned the complexity of the ecological network (i.e., network connectivity) and evaluated the performance of different models at varying levels of the training sample size (**Fig.1C-E**). Here, the network connectivity represents the probability of two species in the species pool interacting with each other. As expected, we observed that the predictive performance of machine learning models improved with the number of training samples. For network connectivity *C* = 0,3, we found that the Area Under the Receiver Operating Characteristic curve (AUROC) of three machine learning models was above 0.9 with training sample size *S*_train_ = *N*. For higher network connectivity (e.g., *C* = 0,4 and 0,5), the increased complexity in inter-species interactions rendered the binary prediction of colonization outcomes more difficult. Nevertheless, with a sample size on the order of ∼*O(N)*), machine learning models were able to achieve accurate classification of colonization outcomes in synthetic data (AUROC > 0.8).

Next, we addressed the *regression* problem, i.e., predicting the steady-state abundance of the exogenous species. For the GLV model, our analytical derivations discovered a surprisingly simple linear relation between the post-invasion abundance of the exogenous species and the pre-invasion abundance of resident species (Supplementary Text, **Fig.S1)**. Although the linear relation doesn’t hold for other dynamical models, it suggests that learning the mapping for COP is feasible by the data-driven approach, and the number of parameters required for fitting the relation is on the order of ∼*O(N*). We employed three machine learning models: Elastic Net Linear Regression, Random Forest regressor, and COP-NODE regressor (**Fig.1F-H**). The predictive performance was evaluated with Pearson’s correlation coefficient between the predicted and true abundance (log-transformed). We systematically examined the predictive performance of three models at varying levels of network connectivity and training sample size. Similar to the classification problem, we found that increasing network connectivity *C* rendered the regression problem more difficult. For training sample size *S*_train_ = 2*N* or higher, there was a substantial improvement in the quantitative prediction of the post-invasion abundance.

### Generation of human stool-derived *in vitro* microbial communities with diverse compositional profiles

To systematically study colonization outcomes in complex microbial communities, we established large-scale cultivation of human stool-derived *in vitro* communities in multi-well plates (*38-41*) (**Fig.2A, Methods**). Briefly, we cultured gut microbial communities derived from 24 donors to reach steady states after five rounds of serial passaging *in vitro*. To increase the diversity in baseline communities, we treated each donor’s sample with 12 antibiotics from different classes (**Table S1**). Overall, we obtained more than 300 baseline communities with substantial variation in the compositional profiles at the species level (**Fig.2B, Fig.S2-S3**). The compositional profiles of the baseline communities were stable, with around 40 to 120 co-existing species in each community (**Fig.S4)**.

**Figure 2.**
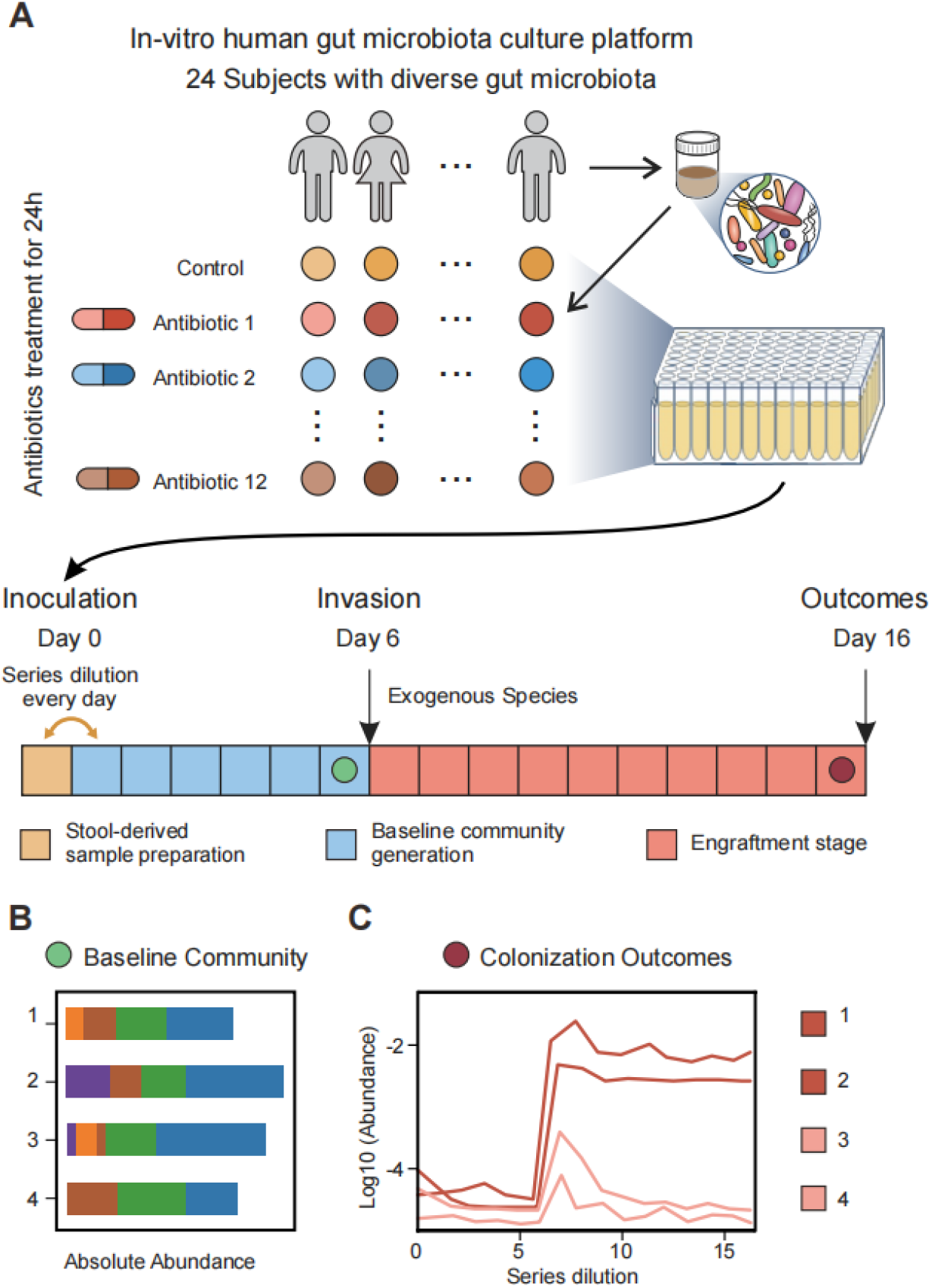
Large-scale invasion experiments in human stool-derived *in vitro* microbial communities. **(A)** Schematic representation of in vitro culture of human stool-derived microbial communities in 96-well plates (**Methods**). Stool samples from 24 donors were treated with 12 different antibiotics for 24 h. The control group was not treated with antibiotics. All communities were passaged five times to reach a stable state, i.e., baseline community profiles (green dot) **(B)**. Exogenous species was introduced on Day 6. After 8-10 times of passaging, the end-point community profiles (red dot) were sequenced to determine the colonization outcome **(C)**.

For the invasion experiments, we would introduce an exogenous species into the baseline communities and determine its colonization outcome after 8-10 rounds of serial passaging (**Fig.2C**). We conducted a preliminary experiment to investigate the colonization outcome of different exogenous species (**Fig.S5**). We found that *E. faecium, A. muciniphila*, and *F. nucleatum* could successfully colonize in some communities at varying levels of post-invasion abundance. In contrast, *S. salivarius, B. breve*, and *Lactobacillus spp*. could not colonize in nearly all the communities we tested. Moreover, vancomycin treatment significantly altered the colonization outcomes, rendering the gut microbial communities more susceptible to invasion (**Fig.S5C**). Overall, our results support the use of human stool-derived *in vitro* communities as a model experimental system for studying colonization outcomes.

### Colonization outcome of *E. faecium* in human stool-derived *in vitro* communities is baseline-dependent and predictable

We selected *E. faecium* as a representative species for colonization experiments in human stool-derived *in vitro* communities. *E. faecium* is a Gram-positive bacterium that inhabits the gut of humans and other animals. Some *E. faecium* strains have probiotic potential (*42*), and recent studies suggest that it plays a positive role in cancer immunotherapy (*43, 44*). On the other hand, some *E. faecium* strains cause opportunistic infections in hospitalized patients with disrupted gut microbiota (*45*).

We introduced *E. faecium* to ∼300 baseline communities (**Fig.S2**) at a dose of 5% relative to the total abundance of resident species. We passaged all communities for ten rounds to reach the post-invasion steady state (**Methods**). We observed that the colonization outcome of *E. faecium* in different communities was persistent during serial passaging (**Fig.S6**). In addition, the composition of *in vitro* communities before and after *E. faecium* invasion is highly reproducible across three replicates (**Fig.S7**).

We found that *E. faecium* was able to colonize 32% of baseline communities (i.e., permissive), with its post-invasion abundance in permissive communities varying over two orders of magnitude (**Fig.3A**). Previous studies suggested that community biomass and diversity are important factors underlying the colonization resistance to exogenous species (*46, 47*). For example, reduced diversity of the resident community is often linked to pathogen infection in the human gut or other ecosystems (*48*). Indeed, we found that the biomass and species richness of the baseline communities exhibited a clear negative correlation with the post-colonization abundances of *E. faecium* (**Fig.S8**). The diversity of the *E. faecium* permissive communities was significantly lower than the resistant communities (**Fig.3B-C**). Furthermore, we observed a significant difference between the composition of *E. faecium* permissive communities and resistant communities (**Fig.3D**). The colonization success of *E. faecium* was highly baseline-dependent, with substantial variations across different donors and antibiotics treatments (**Fig.S9**).

**Figure 3.**
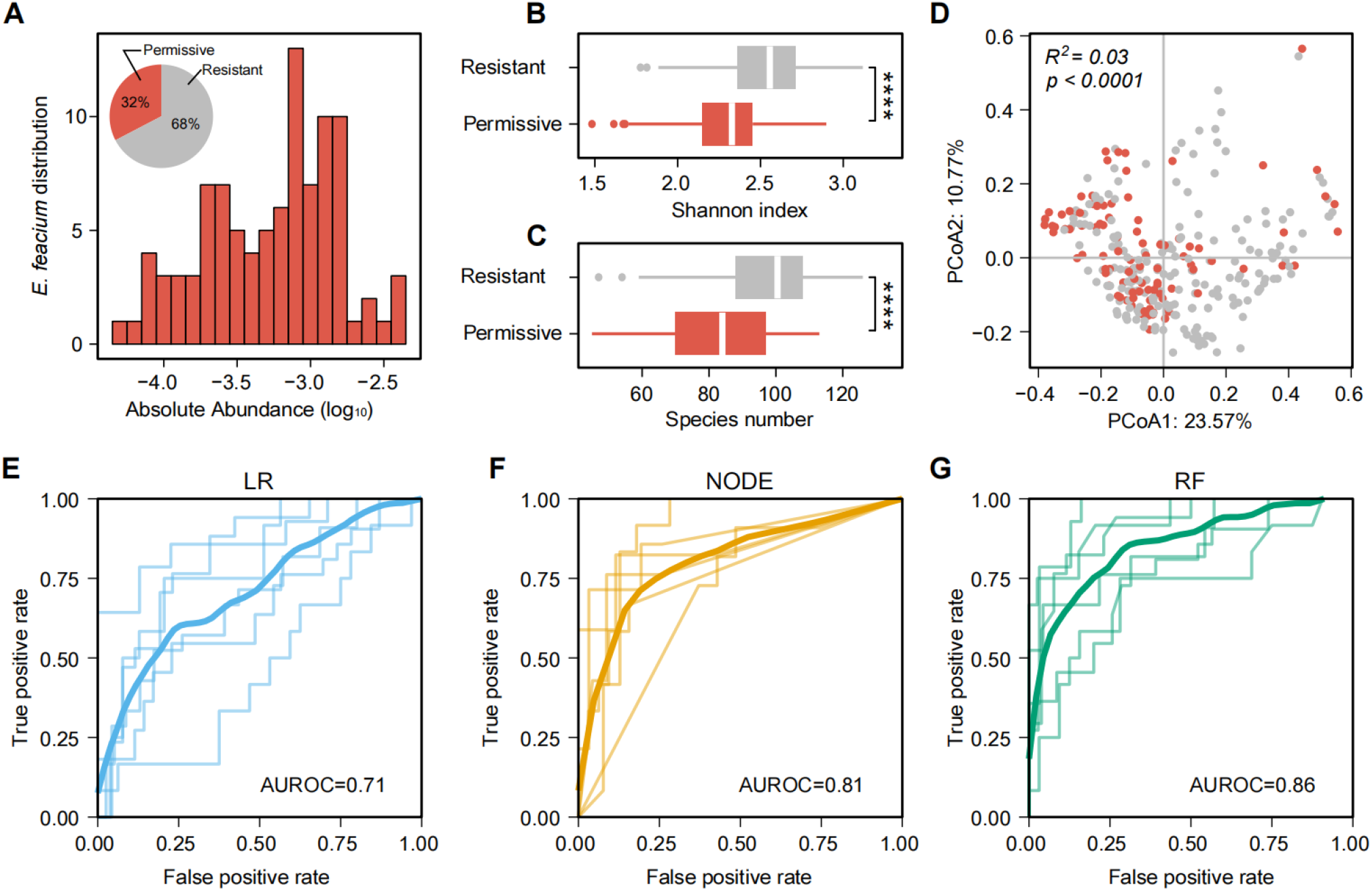
The colonization outcome of *E. faecium* in human stool-derived *in vitro* microbial communities is baseline-dependent and predictable. **(A)** The distribution of *E. faecium* colonization outcomes across permissive communities (colored in red). The abundance of *E. faecium* in resistant communities is below the detection threshold and not shown. **Inset:** Percentage of permissive (red) and resistant communities (gray) based on *E. faecium* colonization outcomes. (**B-C)** Shannon diversity and species richness of *E. faecium* resistant and permissive communities (****p < 0.0001, Mann-Whitney U-tests). **(D)** Principal component analysis (PCoA) based on the Bray-Curtis dissimilarity of the compositional profiles of baseline communities. The difference between permissive and resistant communities was significant (PERMANOVA Adonis test, *R*^2^ = 0,03, *p* < 0,0001). **(E-G)** ROC curve of machine learning models in binary classification (permissive vs. resistant) of the colonization outcomes of *E. faecium*. For each 6-fold cross-validation (ROC curves shown in a light color), we used the samples from 20 subjects to train each model and the samples from the remaining four subjects to evaluate the model. The mean ROC curve is shown in dark color. LR: Logistic Regression, NODE: COP-Neural Ordinary Differential Equations classifier, RF: Random Forest classifier.

To predict the binary colonization outcomes (permissive vs. resistant) of *E. faecium*, we employed three machine learning models, including Logistic Regression, COP-NODE classifier, and Random Forest classifier (**Fig.3E-G, Fig.S10**). For 6-fold cross-validation, we used the communities derived from 20 donors (∼240 samples) to train the model and the communities derived from the remaining 4 donors (∼60 samples) to evaluate the model. Random Forest classifier displayed the best performance in predicting whether *E. faecium* could successfully colonize based on the species-level community composition (AUROC=0.86), followed by COP-NODE classifier (AUROC=0.81) and Logistic Regression (AUROC=0.71). Thus, our colonization experiments of *E. faecium* in complex human gut microbial communities validated that the data-driven approach can solve the classification problem of COP.

### Colonization outcome of *A. muciniphila* is quantitatively predictable

To investigate the generality of our approach, we selected *A. muciniphila* as a second representative species for colonization experiments in human stool-derived *in vitro* communities. *A. muciniphila* is a Gram-negative mucin-degrading bacterium that inhabits the human gut. Due to its potential beneficial effects on human health (*49-51*), *A. muciniphila* is considered a promising probiotic candidate (*52*). *A. muciniphila* is found in the gut microbiome of around 30% of adults, and its abundance varies substantially across individuals (*53*). Similar to the experimental design of *E. faecium*, we introduced *A. muciniphila* to ∼300 baseline communities at a dose of 5% relative to the total abundance of resident species and passaged all communities for eight rounds to reach the post-invasion steady state. The colonization outcome of *A. muciniphila* in different communities was persistent during serial passaging (**Fig.S11**), and the composition of *in vitro* communities post *A. muciniphila* invasion is highly reproducible across three replicates (**Fig.S12**).

Overall, we found substantial variations in the post-invasion steady-state abundance of *A. muciniphila* across different donors and antibiotics treatments (**Fig.S13**). *A. muciniphila* could colonize in 93.6% of baseline communities (i.e., permissive). For permissive communities, the post-invasion abundance of *A. muciniphila* displayed a bimodal distribution (**Fig.4A**). We classified the permissive communities into two subgroups (high vs. low), depending on the post-invasion abundance of *A. muciniphila* (abundance threshold at 10^−2^). The Shannon diversity (**Fig.4B**) and species richness (**Fig.4C**) of the *A. muciniphila* high permissive communities were significantly lower than those of the low permissive communities, and there was a significant difference between the community composition of the two groups (**Fig.4D**).

**Figure 4.**
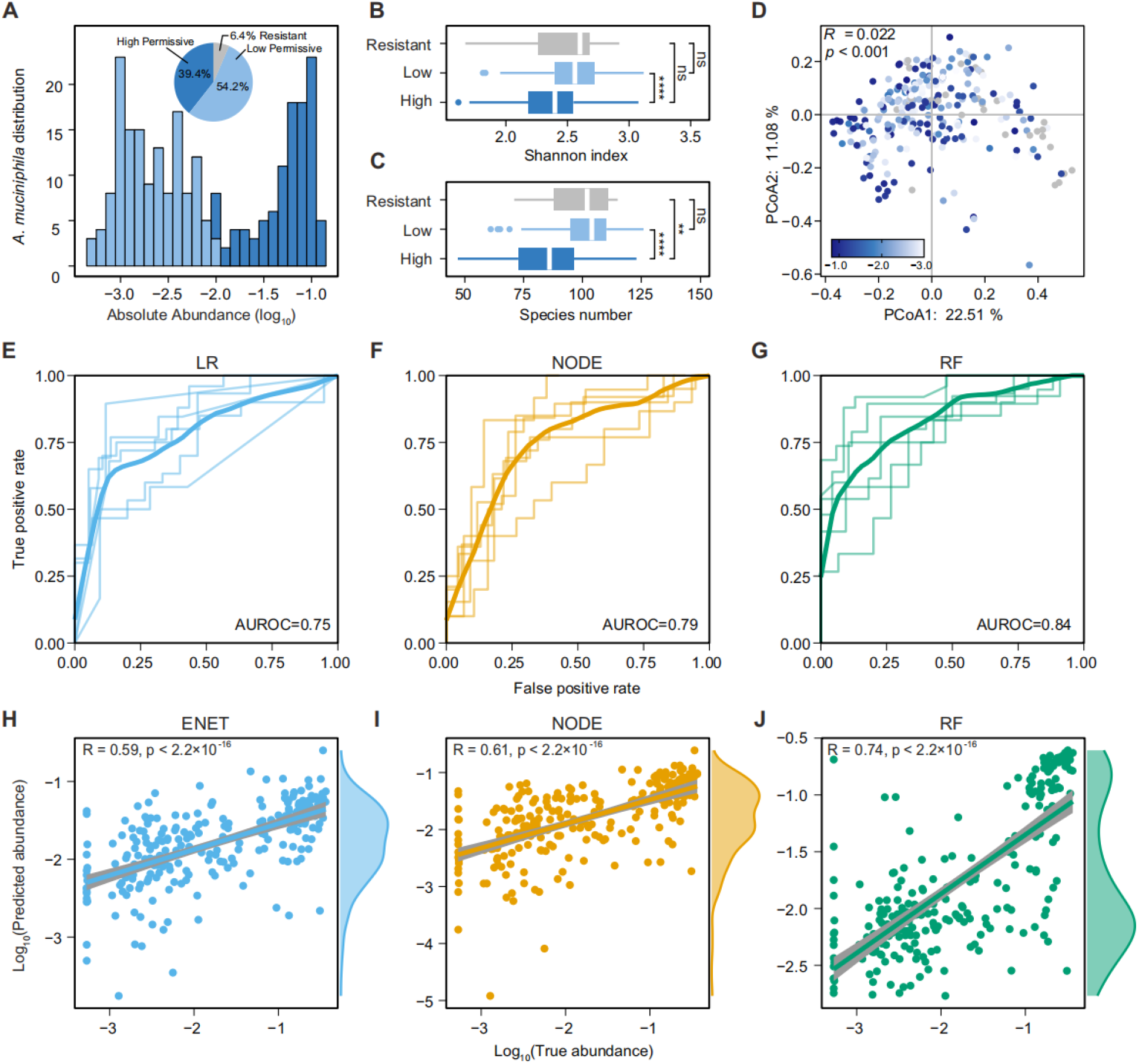
The colonization outcome of *A. muciniphila* in human stool-derived *in vitro* microbial communities is quantitatively predictable. **(A)** The distribution of *A. muciniphila* colonization outcomes across permissive communities. The abundance of *A. muciniphila* in resistant communities is below the detection threshold and not shown. **Inset:** Percentage of high permissive (dark blue color), low permissive (light blue), and resistant communities (gray) based on *A. muciniphila* colonization outcomes. **(B-C)** Shannon diversity and species richness of *A. muciniphila* resistant, permissive (low), and permissive (high) communities (ns, not significant, **p < 0.01,****p < 0.0001, Mann-Whitney U-tests). **(D)** Principal component analysis (PCoA) plots based on the Bray-Curtis dissimilarity of the compositional profiles of baseline communities. Color of the point showing the abundance of *A. muciniphila* in communities. The difference between highly permissive and lowly permissive communities was significant (PERMANOVA Adonis test, *R*^2^ = 0,022, *p* < 0,001). **(E-G)** ROC curve of machine learning models in binary classification (high permissive vs. low permissive) of the colonization outcomes of *A. muciniphila*. For each 6-fold cross-validation (ROC curves shown in a light color), we used the samples from 20 subjects to train each model and the samples from the remaining four subjects to evaluate the model. The mean ROC curve is shown in dark color. ENET: Elastic Net Linear Regression, NODE: COP-Neural Ordinary Differential Equations regressor, RF: Random Forest regressor. **(H-J)** Pearson’s correlation coefficient between the predicted abundance and the true abundance of *A. muciniphila*.

We evaluated the performance of machine learning models in predicting the colonization outcome of *A. muciniphila*, both qualitatively (classification) and quantitatively (regression). Random Forest classifier displayed the best performance in binary classification (high permissive vs. low permissive of *A. muciniphila*) based on the species-level community composition (AUROC=0.84), followed by COP-NODE classifier (AUROC=0.79) and Logistic Regression (AUROC=0.74) (**Fig.4E-G**). To quantitatively predict the post-invasion abundance of *A. muciniphila*, we employed three machine learning models: Elastic Net Linear Regression, COP-NODE regressor, and Random Forest regressor (**Fig.4H-J, Fig.S14**). In comparison to the other two methods, the Random Forest regressor achieved the highest accuracy in quantitative prediction (Pearson’s correlation coefficient between the predicted and true abundances R = 0,74, *p* < 2,2 × 10^−16^) and successfully recapitulated the bimodal distribution in the abundance of *A. muciniphila*. Taken together, we demonstrated the generality of the data-driven approach in predicting baseline-dependent colonization outcomes for complex microbial communities.

### Colonization impact in simulated and experimental communities

Learning the mapping from the baseline taxonomic profile to colonization outcomes can help us infer the impact of each resident species on the colonization of the exogenous species (**Fig.1B**). To compute the colonization impact of regression (classification), we can perform a thought experiment by introducing a perturbation in the abundance of the resident species, and use the trained machine learning model to predict the new colonization outcome of invading species after the perturbation (**Fig.5A**). Negative colonization impact means that a resident species inhibits the colonization of the exogenous species in a given local community. In GLV simulations, while the colonization impact of resident species was randomly distributed (**Fig.5B**), we found that the median colonization impact of a resident species across different local communities was positively correlated to its interaction strength on the exogenous species (Spearman correlation coefficient *ρ* = 0,73, *p* < 2,2 × 10^−16^, **Fig.5C**), suggesting that we may use colonization impact to identify strongly interacting species.

**Figure 5.**
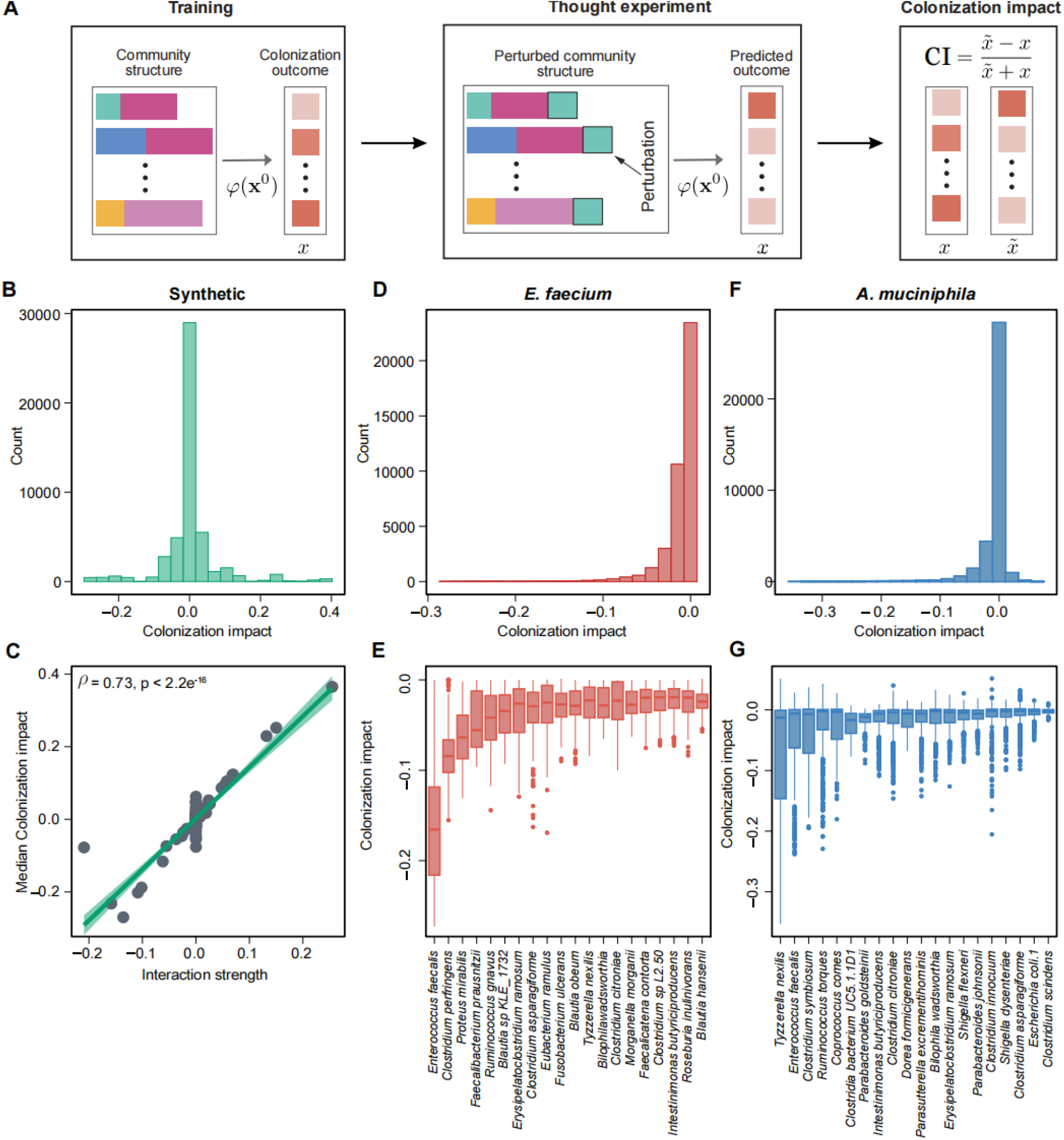
Colonization impact in simulated and experimental communities. **(A)** To compute the colonization impact, i.e., the impact of a resident species on the colonization outcome of the invading species, we first trained the prediction models using all the samples. Then, we performed a thought experiment by introducing a perturbation in the abundance of the resident species and used the trained machine learning model to predict the new steady state abundance of invading species after the perturbation. A negative colonization impact indicates that a resident species inhibits the colonization of the invading species. **(B-C)** In simulated data, the colonization impact is randomly distributed. The median colonization impact of a resident species across different local communities is positively correlated to its interaction strength on the exogenous species (Spearman correlation *ρ* = 0,73, *p* < 2,2 × 10^−16^),). Network connectivity *C* = 0,3, *S*_trian_/*N* = 5. COP-NODE regressor is used. **(D-E)** The distribution of colonization impact on *E. faecium*, and the top ranking species with negative colonization impact (median across different communities, RF classifier). **(F-G)** The distribution of colonization impact on *A. muciniphila* and the top-ranking species with negative colonization impact (median across different communities, RF regressor).

We used the Random Forest model to evaluate the colonization impact of all species in human stool-derived *in vitro* communities on *E. faecium* and *A. muciniphila* (**Fig.5D-G**). We inferred that most resident species had a weak negative colonization impact (**Fig.5D and 5F**). Based on the median colonization impact of a certain resident species across different local communities, we identified the top-ranking species with negative colonization impact (**Fig.5E and 5G**). Colonization impact on *A. muciniphila* was overall less negative than *E. faecium*, consistent with our observation that human gut microbial communities were more permissive to *A. muciniphila* colonization.

### The impact of strongly interacting species on colonization outcomes

To understand the role of strongly interacting species on colonization outcomes, we systematically studied the impact of *E. faecalis* on the colonization of *E. faecium. E. faecalis* was inferred to have the strongest colonization impact on *E. faecium* across different baseline communities (**Fig.5E**). We observed a statistically significant negative correlation between the abundance of *E. faecalis* in baseline communities and the post-invasion abundance of *E. faecium* (Kendall’s τ = −0,37, *p* = 5,29 × 10^−16^). In particular, baseline communities derived from some donors (e.g., S10, S07) had a high abundance of *E. faecalis* and were resistant to *E. faecium* colonization (**Fig.6A)**.

**Figure 6.**
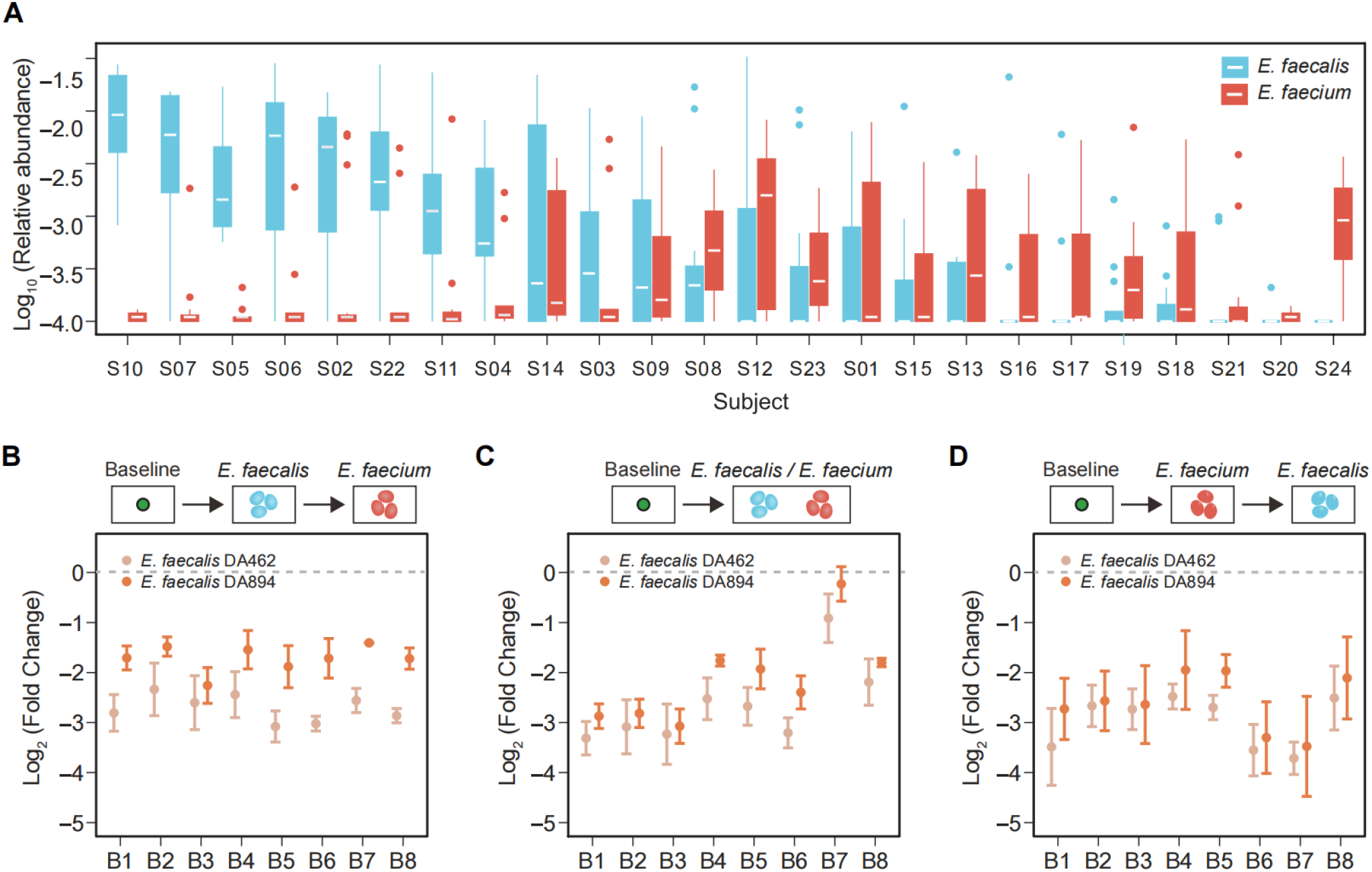
The presence of *E. faecalis* in baseline communities inhibits the invasion of *E. faecium*. **(A)** The post-invasion relative abundance of *E. faecium* (aqua) is negatively associated with the relative abundance of *E. faecalis* (red) across baseline communities derived from different human subjects (labeled as S01 to S24). **(B-D)** The colonization of *E. faecium* is significantly inhibited by *E. faecalis* across different baseline communities (labeled as B1 to B8). There were three different intervention groups: 1) add *E. faecalis* (or *C. symbiosum*) into the baseline community, followed by *E. faecium* on the next day (**B**); 2) add *E. faecalis* and *E. faecium* on the same day (**C**); 3) add *E. faecium* into the baseline community, followed by *E. faecalis* on the next day **(D**). In the control group, we only added *E. faecium*. After five passages, the end-point abundance of *E. faecium* was measured by qPCR. The fold change in the end-point abundance of *E. faecium* (the intervention group divided by the control group) is lower than 1 (dashed line), indicating that *E. faecalis* inhibits the colonization of *E. faecium*. Two different *E. faecalis* strains, DA462 and DA894, were used. n= 3 replicates, the error bars are SEMs.

We found that *E. faecalis* inhibited the growth of *E. faecium* in pairwise co-culture, either in liquid culture or on agar plates (**Fig.S15**). Then, we introduced *E. faecalis* into eight human stool-derived *in vitro* communities that were permissive to *E. faecium* invasion, using three different types of interventions (**Fig.6B-D, Fig.S16A**): 1) add *E. faecalis* into the baseline community, followed by *E. faecium* on the next day; 2) add *E. faecalis* and *E. faecium* on the same day; 3) add *E. faecium* into the baseline community, followed by *E. faecalis* on the next day. In the control group, we only added *E. faecium*. In all three intervention groups, the colonization of *E. faecium* was significantly inhibited by *E. faecalis* across different baseline communities. Also, the inhibitory effect was consistent for two different *E. faecalis* strains isolated from human stool samples (**Methods**). In comparison, *Clostridium symbiosum*, a species predicted to have a neutral impact, did not alter the colonization of *E. faecium* (**Fig.S16B**).

Finally, we explored if the strong inhibition of *E. faecalis* on *E. faecium* could be shaping their distribution in the human gut via priority effects, i.e., the gut microbiome colonized with *E. faecalis* becomes resistant to *E. faecium*. We performed metagenomic sequencing of ∼120 healthy volunteers in the SIAT cohort (**Methods**), whose samples were used to derive the *in vitro* communities and isolate the *Enterococcus* strains in this study. Indeed, there was a statistically significant negative correlation between the relative abundance of *E. faecalis* and *E. faecium* in the SIAT cohort (Kendall correlation τ = −0,36, *p* = 0,0044, **Fig. S17A**). A similar pattern was observed in gut metagenomic samples of four independent cohorts (Kendall correlation τ = −0,36, *p* = 5,439 × 10^−15^), **Fig. S17B**).

Overall, our experimental validations and analysis suggest that data-driven models can infer species with strong colonization impact and guide the modulation of resident communities to alter the colonization outcomes of exogenous species.

## Discussion

Here we proposed and systematically validated a data-driven approach to predict colonization outcomes of exogenous species, providing a powerful tool to inform the management of complex ecosystems. Pairwise co-culture (*47, 54-56*) and synthetic communities (*17, 57, 58*) have been widely used to study the ecology and function of microbial communities. These experiments require the isolation and cultivation of individual species, thus are often limited to simple communities. In comparison, our approach is based on sampling an ensemble of complex communities (∼100 species, **Fig.S4**) and using the sampled communities to infer the mapping between community composition and colonization outcomes (*59*). We demonstrate that the data-driven approach enables accurate function prediction and system-level understanding of complex microbial communities.

Understanding the colonization resistance of complex communities is a fundamental question in ecology. In our large-scale invasion experiments (∼300 local communities and two different exogenous species), we found that resistance to exogenous species was positively correlated to community diversity, supporting the view that colonization resistance is an emergent property of complex communities (*8*). While most resident species had a weak negative impact on the colonization of exogenous species, we identified *E. faecalis* as a strong inhibitor of *E. faecium*. We validated that introducing strongly interacting species into baseline communities can alter the colonization outcomes. It should be noted that the colonization impact is dependent on the community context (**Fig.5**), because it takes into account both direct and indirect effects on the invading species (*60*), as well as potential higher-order interactions (*61, 62*). Previous studies have shown that strongly interacting species can lead to priority effects (*63*), with important implications for community assembly in the infant gut microbiome and the formation of community types (*64, 65*). Moreover, strongly interacting species can be used to modulate the resident communities to prevent the colonization of pathogens (*16*) or facilitate the colonization of beneficial microbes (e.g., probiotics, crop fertilizers) (*66*).

Our results suggest that the colonization resistance of microbial communities is predictable and tunable via the data-driven approach, given that training data size is sufficient (on the order of ∼*O(N)*). The high-throughput cultivation of gut microbial communities *in vitro* provides a powerful approach to studying the human gut microbiome (*67, 68*). In our experiments, the number of species in the meta-community was ∼160, and we profiled ∼300 baseline communities for proof-of-concept validation. Meeting the sample size requirement for gnotobiotic plants is feasible (*69, 70*). However, it could be challenging to gather sufficient training data for gnotobiotic animals and human cohort studies, depending on the complexity of the meta-community. In addition to data size, another critical concern is the technical variability in large-scale experiments (*71*). In future studies, experimental workflows can be automated to minimize technical variability and ensure data quality for training machine learning models.

Our study has several limitations. First, we did not account for potential variations at the strain level (*72*). Previous studies have shown that the strength of interspecies interactions can vary across different strains, such as the inhibition of *Klebsiella pneumoniae* by *Klebsiella oxytoca (16, 73*). We also observed strain-level variations in the inhibition of *E. faecium* by *E. faecalis* (**Fig.6**), and the underlying mechanism remains to be elucidated. Second, our invasion experiments *in vitro* did not reflect host-mediated interactions, which also contribute to colonization resistance *in vivo (22*). Nevertheless, the higher permissiveness to *A. muciniphila* than *E. faecium* in human gut microbial communities *in vitro* is consistent with the higher prevalence of *A. muciniphila* in metagenomic samples (*53, 74*). Third, we assumed that there was a single post-invasion steady state in simulated and experimental communities. The colonization outcomes may be influenced by multi-stability in microbial communities, e.g., successful colonization depends on the initial abundance of the invading species (*75, 76*).

Our data-driven approach is independent of any dynamics model to predict colonization outcomes of exogenous species for complex microbial communities without detailed knowledge of the underlying ecological and biochemical process. We anticipate that the data-driven approach can be generalized to predict and engineer the function of microbial communities (i.e. mapping from community composition to function) (*36, 77-79*). Similarly, this approach can be used to predict the response of microbial communities to various types of perturbations (i.e. mapping from community composition to the shift in composition/function), such as the baseline-dependent response of the human gut microbiome to prebiotics, food additives, etc. (*80, 81*). In parallel to the breakthroughs in predicting the properties of complex biomolecules, we envision that the data-driven approach will lead to a paradigm shift in studying the stability and function of complex ecological systems and guide important applications in healthcare (e.g. personalized nutrition based on the human gut microbiome) and agriculture.

## Acknowledgments

We thank Na Li and volunteers at Shenzhen Institute of Advanced Technology (SIAT) for stool sample collection. We thank Zepeng Qu and Zhenkun Zhang for isolating human gut bacterial strains. We thank Prof. Jiachao Zhang at Hainan University for providing probiotic strains. We thank Huaijie Hao and Yan Tan at Xbiome for providing help with the anaerobic workstation. We thank Shenzhen Infrastructure for Synthetic Biology and Chaobi Lei for providing support in DNA extraction. We thank Zheng Sun, Chen Liao, Hongbin Liu, Lanxiang Wang, and colleagues at SIAT for valuable discussions.

## Author contributions

L.D., Y.Y.L., L.W., and X.W.W. conceived the presented idea, designed the project, interpreted the results, and wrote the manuscript. X.W.W. implemented in silico simulations, machine learning models, and analytical derivations. L.W. and Z.T. performed the in vitro experiments. Y.Z. assisted in experiments. L.W. analyzed the experimental data. T.W. discussed the results and revised the manuscript. W.Z. assisted in data analysis and interpretation. All authors approved the final manuscript. L.D. and Y.Y.L. provided supervision and resources for this study.

## Data availability

All sequencing data generated in this study are available from European Nucleotide Archive (ENA) under study accession number PRJEB60398.

## Code availability

The code for simulations and data analysis is available at https://github.com/spxuw/COP.

## References

1. C. G. Buffie, E. G. Pamer, Microbiota-mediated colonization resistance against intestinal pathogens. Nature reviews. Immunology 13, 790–801 (2013).

2. T. Gensollen, S. S. Iyer, D. L. Kasper, R. S. Blumberg, How colonization by microbiota in early life shapes the immune system. Science 352, 539–544 (2016).

3. J. Walter, M. X. Maldonado-Gomez, I. Martinez, To engraft or not to engraft: an ecological framework for gut microbiome modulation with live microbes. Curr Opin Biotechnol 49, 129–139 (2018).

4. D. R. Amor, C. Ratzke, J. Gore, Transient invaders can induce shifts between alternative stable states of microbial communities. Sci. Adv. 6, (2020).

5. N. I. van den Berg et al., Ecological modelling approaches for predicting emergent properties in microbial communities. Nat Ecol Evol 6, 855–865 (2022).

6. T. A. Kennedy et al., Biodiversity as a barrier to ecological invasion. Nature 417, 636–638 (2002).

7. H. M. Kurkjian, M. J. Akbari, B. Momeni, The impact of interactions on invasion and colonization resistance in microbial communities. PLoS Comput Biol 17, e1008643 (2021).

8. T. J. Case, Invasion resistance arises in strongly interacting species-rich model competition communities. Proc Natl Acad Sci U S A 87, 9610–9614 (1990).

9. P. Vonaesch, M. Anderson, P. J. Sansonetti, Pathogens, microbiome and the host: emergence of the ecological Koch’s postulates. FEMS Microbiol Rev 42, 273–292 (2018).

10. E. G. Pamer, Resurrecting the intestinal microbiota to combat antibiotic-resistant pathogens. Science 352, 535–538 (2016).

11. P. D. Li et al., The phyllosphere microbiome shifts toward combating melanose pathogen. Microbiome 10, 56 (2022).

12. R. Mendes, P. Garbeva, J. M. Raaijmakers, The rhizosphere microbiome: significance of plant beneficial, plant pathogenic, and human pathogenic microorganisms. FEMS Microbiol Rev 37, 634–663 (2013).

13. T. S. Schmidt et al., Extensive transmission of microbes along the gastrointestinal tract. eLife 8, (2019).

14. C. R. Kok, R. Hutkins, Yogurt and other fermented foods as sources of health-promoting bacteria. Nutr Rev 76, 4–15 (2018).

15. B. Chassaing, E. Cascales, Antibacterial Weapons: Targeted Destruction in the Microbiota. Trends Microbiol 26, 329–338 (2018).

16. S. G. Kim et al., Microbiota-derived lantibiotic restores resistance against vancomycin-resistant Enterococcus. Nature 572, 665–669 (2019).

17. C. Eberl et al., E. coli enhance colonization resistance against Salmonella Typhimurium by competing for galactitol, a context-dependent limiting carbon source. Cell Host & Microbe 29, 1680–1692 e1687 (2021).

18. R. A. Oliveira et al., Klebsiella michiganensis transmission enhances resistance to Enterobacteriaceae gut invasion by nutrition competition. Nat Microbiol 5, 630–641 (2020).

19. Y. Litvak et al., Commensal Enterobacteriaceae Protect against Salmonella Colonization through Oxygen Competition. Cell Host & Microbe 25, 128-139.e125 (2019).

20. D. P. Lloyd, R. J. Allen, Competition for space during bacterial colonization of a surface. Journal of the Royal Society Interface 12, (2015).

21. Ivanov, II et al., Induction of intestinal Th17 cells by segmented filamentous bacteria. Cell 139, 485–498 (2009).

22. B. Stecher et al., Salmonella enterica serovar typhimurium exploits inflammation to compete with the intestinal microbiota. PLoS Biol 5, 2177–2189 (2007).

23. C. Human Microbiome Project, Structure, function and diversity of the healthy human microbiome. Nature 486, 207–214 (2012).

24. E. A. Franzosa et al., Identifying personal microbiomes using metagenomic codes. Proc Natl Acad Sci U S A 112, E2930–2938 (2015).

25. N. Han et al., Time-scale analysis of the long-term variability of human gut microbiota characteristics in Chinese individuals. Commun Biol 5, 1414 (2022).

26. P. W. O’Toole, I. B. Jeffery, Gut microbiota and aging. Science 350, 1214–1215 (2015).

27. P. J. Turnbaugh, F. Backhed, L. Fulton, J. I. Gordon, Diet-induced obesity is linked to marked but reversible alterations in the mouse distal gut microbiome. Cell Host & Microbe 3, 213–223 (2008).

28. L. Maier et al., Extensive impact of non-antibiotic drugs on human gut bacteria. Nature 555, 623-+ (2018).

29. S. Alavi et al., Interpersonal Gut Microbiome Variation Drives Susceptibility and Resistance to Cholera Infection. Cell 181, 1533-+ (2020).

30. C. G. Buffie et al., Precision microbiome reconstitution restores bile acid mediated resistance to Clostridium difficile. Nature 517, 205–U207 (2015).

31. M. X. Maldonado-Gomez et al., Stable Engraftment of Bifidobacterium longum AH1206 in the Human Gut Depends on Individualized Features of the Resident Microbiome. Cell Host & Microbe 20, 515–526 (2016).

32. N. Zmora et al., Personalized Gut Mucosal Colonization Resistance to Empiric Probiotics Is Associated with Unique Host and Microbiome Features. Cell 174, 1388–1405 e1321 (2018).

33. C. Liao et al., Compilation of longitudinal microbiota data and hospitalome from hematopoietic cell transplantation patients. Sci Data 8, 71 (2021).

34. J. Jumper et al., Highly accurate protein structure prediction with AlphaFold. Nature 596, 583–589 (2021).

35. E. D. Vaishnav et al., The evolution, evolvability and engineering of gene regulatory DNA. Nature 603, 455–463 (2022).

36. S. Michel-Mata, X. W. Wang, Y. Y. Liu, M. T. Angulo, Predicting microbiome compositions from species assemblages through deep learning. iMeta 1, (2022).

37. A. Bashan et al., Universality of human microbial dynamics. Nature 534, 259–262 (2016).

38. E. Hernandez-Sanabria, J. F. Vazquez-Castellanos, J. Raes, In vitro ecology: a discovery engine for microbiome therapies. Nat Rev Gastroenterol Hepatol, (2020).

39. B. Javdan et al., Personalized Mapping of Drug Metabolism by the Human Gut Microbiome. Cell, (2020).

40. L. Li et al., RapidAIM: a culture- and metaproteomics-based Rapid Assay of Individual Microbiome responses to drugs. Microbiome 8, 33 (2020).

41. A. Aranda-Diaz et al., Establishment and characterization of stable, diverse, fecal-derived in vitro microbial communities that model the intestinal microbiota. Cell Host & Microbe 30, 260–272 e265 (2022).

42. H. Hanchi, W. Mottawea, K. Sebei, R. Hammami, The Genus Enterococcus: Between Probiotic Potential and Safety Concerns-An Update. Front Microbiol 9, 1791 (2018).

43. M. E. Griffin et al., Enterococcus peptidoglycan remodeling promotes checkpoint inhibitor cancer immunotherapy. Science 373, 1040–1046 (2021).

44. X. Xiong et al., Emerging enterococcus pore-forming toxins with MHC/HLA-I as receptors. Cell 185, 1157–1171 e1122 (2022).

45. D. Van Tyne, M. S. Gilmore, Friend turned foe: evolution of enterococcal virulence and antibiotic resistance. Annu Rev Microbiol 68, 337–356 (2014).

46. M. L. Jones, D. W. Rivett, A. Pascual-Garcia, T. Bell, Relationships between community composition, productivity and invasion resistance in semi-natural bacterial microcosms. eLife 10, (2021).

47. S. Hromada et al., Negative interactions determine Clostridioides difficile growth in synthetic human gut communities. Mol Syst Biol 17, e10355 (2021).

48. F. Keesing et al., Impacts of biodiversity on the emergence and transmission of infectious diseases. Nature 468, 647–652 (2010).

49. H. J. You et al., Bacteroides vulgatus SNUG 40005 restores Akkermansia depletion by metabolite modulation. Gastroenterology, (2022).

50. L. Derosa et al., Intestinal Akkermansia muciniphila predicts clinical response to PD-1 blockade in patients with advanced non-small-cell lung cancer. Nat Med 28, 315–324 (2022).

51. M. C. Dao et al., Akkermansia muciniphila and improved metabolic health during a dietary intervention in obesity: relationship with gut microbiome richness and ecology. Gut 65, 426–436 (2016).

52. Q. X. Zhai, S. S. Feng, N. Arjan, W. Chen, A next generation probiotic, Akkermansia muciniphila. Crit Rev Food Sci 59, 3227–3236 (2019).

53. N. Karcher et al., Genomic diversity and ecology of human-associated Akkermansia species in the gut microbiome revealed by extensive metagenomic assembly. Genome Biol 22, 209 (2021).

54. J. Chen et al., A commensal-encoded genotoxin drives restriction of Vibrio cholerae colonization and host gut microbiome remodeling. Proc Natl Acad Sci U S A 119, e2121180119 (2022).

55. A. S. Weiss et al., In vitro interaction network of a synthetic gut bacterial community. ISME J 16, 1095–1109 (2022).

56. F. Getzke et al., Cofunctioning of bacterial exometabolites drives root microbiota establishment. Proc Natl Acad Sci U S A 120, e2221508120 (2023).

57. S. Brugiroux et al., Genome-guided design of a defined mouse microbiota that confers colonization resistance against Salmonella enterica serovar Typhimurium. Nat Microbiol 2, 16215 (2016).

58. J. Kehe et al., Massively parallel screening of synthetic microbial communities. Proc Natl Acad Sci U S A 116, 12804–12809 (2019).

59. C. Gopalakrishnappa, K. Gowda, K. H. Prabhakara, S. Kuehn, An ensemble approach to the structure-function problem in microbial communities. iScience 25, 103761 (2022).

60. Y. Xiao, M. T. Angulo, S. Lao, S. T. Weiss, Y. Y. Liu, An ecological framework to understand the efficacy of fecal microbiota transplantation. Nat Commun 11, 3329 (2020).

61. M. M. Mayfield, D. B. Stouffer, Higher-order interactions capture unexplained complexity in diverse communities. Nat Ecol Evol 1, 62 (2017).

62. A. Sanchez-Gorostiaga, D. Bajic, M. L. Osborne, J. F. Poyatos, A. Sanchez, High-order interactions distort the functional landscape of microbial consortia. PLoS Biol 17, (2019).

63. R. Debray et al., Priority effects in microbiome assembly. Nat Rev Microbiol 20, 109–121 (2022).

64. C. Rao et al., Multi-kingdom ecological drivers of microbiota assembly in preterm infants. Nature 591, 633–638 (2021).

65. T. E. Gibson, A. Bashan, H.-T. Cao, S. T. Weiss, Y.-Y. Liu, On the Origins and Control of Community Types in the Human Microbiome. PLoS Comput Biol 12, (2016).

66. H. J. You et al., Bacteroides vulgatus SNUG 40005 Restores Akkermansia Depletion by Metabolite Modulation. Gastroenterology 164, 103–116 (2023).

67. G. Vrancken, A. C. Gregory, G. R. B. Huys, K. Faust, J. Raes, Synthetic ecology of the human gut microbiota. Nat Rev Microbiol 17, 754–763 (2019).

68. A. Aranda-Diaz et al., Establishment and characterization of stable, diverse, fecal-derived in vitro microbial communities that model the intestinal microbiota. Cell Host & Microbe 30, 260–272 e265 (2022).

69. C. M. Vogel, D. B. Potthoff, M. Schafer, N. Barandun, J. A. Vorholt, Protective role of the Arabidopsis leaf microbiota against a bacterial pathogen. Nat Microbiol 6, 1537–1548 (2021).

70. S. Herrera Paredes et al., Design of synthetic bacterial communities for predictable plant phenotypes. PLoS Biol 16, e2003962 (2018).

71. C. van de Velde et al., Technical versus biological variability in a synthetic human gut community. Gut Microbes 15, 2155019 (2023).

72. H. Hu et al., StrainPanDA: Linked reconstruction of strain composition and gene content profiles via pangenome-based decomposition of metagenomic data. iMeta 1, (2022).

73. L. Osbelt et al., Klebsiella oxytoca causes colonization resistance against multidrug-resistant K. pneumoniae in the gut via cooperative carbohydrate competition. Cell Host & Microbe 29, 1663–1679 e1667 (2021).

74. S. Wu et al., GMrepo: a database of curated and consistently annotated human gut metagenomes. Nucleic Acids Res 48, D545–D553 (2020).

75. D. Gonze, L. Lahti, J. Raes, K. Faust, Multi-stability and the origin of microbial community types. ISME J 11, 2159–2166 (2017).

76. L. Dai, D. Vorselen, K. S. Korolev, J. Gore, Generic Indicators for Loss of Resilience Before a Tipping Point Leading to Population Collapse. Science 336, 1175–1177 (2012).

77. A. M. Schubert, H. Sinani, P. D. Schloss, Antibiotic-Induced Alterations of the Murine Gut Microbiota and Subsequent Effects on Colonization Resistance against Clostridium difficile. mBio 6, e00974 (2015).

78. A. Sanchez et al., The community-function landscape of microbial consortia. Cell Syst 14, 122–134 (2023).

79. R. L. Clark et al., Design of synthetic human gut microbiome assembly and butyrate production. Nat Commun 12, 3254 (2021).

80. N. T. Baxter et al., Dynamics of Human Gut Microbiota and Short-Chain Fatty Acids in Response to Dietary Interventions with Three Fermentable Fibers. mBio 10, (2019).

81. J. Suez et al., Personalized microbiome-driven effects of non-nutritive sweeteners on human glucose tolerance. Cell 185, 3307–3328 e3319 (2022).

