## Supplementary Materials for "Data-driven prediction of colonization outcomes for complex microbial communities"

**The PDF file includes:**

Materials and Methods
Supplementary Text
Figs. S1 to S18
Table S1
References

### Materials and Methods

#### *In silico* simulations of colonization outcomes

We generated synthetic data of colonization outcomes using the generalized Lotka–Volterra (GLV) model (*1*):

$$\frac{dx_i(t)}{dt} = x_i(t)\left[r_i + \sum_{j=1}^{N} a_{ij}x_j(t)\right], i = 1, \cdots, N. \quad (1)$$

Here $x_i(t)$ represents the absolute abundance of the *i*-th species at time $t \geq 0$. The pair-wise microbial interaction is presented by the matrix $A = (a_{ij}) \in \mathbb{R}^{N\times N}$, with $a_{ij} > 0$ ($< 0$, or $= 0$) means that species-*j* promotes (inhibits or does not affect) the growth of species-*i*, respectively. The ecological network $G(A)$ is constructed using an Erdős-Rényi random graph model (*2*) with $N$ nodes (i.e., species) and connectivity $C$ (i.e., the probability connecting two species). To generate the interaction matrix $A$ of ecological network, for each link $(j \rightarrow i) \in G(A)$ with $j \neq i$, we draw $a_{ij}$ from the normal distribution $\mathbb{N}(0, \sigma)$. All other entries of $A$ are set to be zero. The intrinsic growth rate vector $r = [r_i] \in \mathbb{R}^N$ is drawn from a uniform distribution $\mathcal{U}(0,1)$. Each local community includes $N_s$ species randomly drawn from the $(N - 1)$ species (excluding the exogenous species) and $N_s$=30 in all simulations.

To examine the performance of colonization outcome prediction in communities with varying levels of network complexity, we tuned the network connectivity $C$ from the set [0.3, 0.4, 0.5]. In addition, to evaluate the sample size required for accurate prediction, we systematically tuned the size of training samples $S_{\text{train}}/N$ from 0.5 to 10. An independently generated set of 100 samples were used as test data to evaluate the models. To generate the training samples for classification, we selected 1,100 local communities where the post-invasion steady-state abundance of the exogenous species is above 0.05 (i.e., the threshold used to determine successful colonization) in half of the local communities, and below 0.05 in the other half. To generate the training samples for regression, we selected 1,100 local communities in which the post-invasion steady-state abundance of the exogenous species follows the log-normal distribution (mean=-3, standard deviation=0.5).

#### Colonization outcome prediction by machine learning models

We developed a deep learning model for Colonization Outcome Prediction using the Neural Ordinary Differential Equations (COP-NODE) (*3*). The architecture of COP-NODE consists of two fully collected layers, and each fully connected layer (with dimension $N$) is followed by a normalization layer and a ReLU activation layer. The final layer is Sigmoid activation. The Adam optimizer was used for the optimization with a learning rate 0.01 for both classification and regression. The loss function is CrossEntropy for classification and SmoothL1Loss for regression

(*4*). We randomly selected 20% of training samples as the validation set to select the best model and hyperparameters. For classification, we tuned the batch size from the set [16, 32, 64] and the hyperparameter β (the threshold to change between L1 and L2 regularization) from the set [0.001, 0.01, 0.1, 0.2, 0.4, 0.6, 0.8, 1]. Other machine learning models used in this study, including Logistic Regression, Elastic Net, Random Forest classifier, and regressor, were implemented using the Python package scikit-sklearn (*5*). We used randomized search on hyperparameters and 3-fold cross-validation to optimize the AUROC for classification and $R^2$ for regression. The regression models were trained to predict the log-transformed abundance of the exogenous species.

**Collection and preservation of human stool samples**

Stool samples were collected from healthy human donors and were immediately transferred to an anaerobic workstation (85% $N_2$, 10% $H_2$ and 5% $CO_2$, COY). 10g of each stool sample was suspended into 50mL 20% glycerol (v/v, in sterile phosphate-buffered saline, with 0.1% L-cysteine hydrochloride), homogenized by vortexing, and then filtered with sterile nylon mesh to remove large particles in fecal matter. Aliquots of the suspension were stored in sterile cryogenic vials and frozen at -80 °C for long-term storage until processing for DNA extraction and culturing so that the stool-derived community could be revived (thawed) for repeatable experiments. The collection of human stool samples from volunteers at SIAT (referred to as "SIAT cohort") were approved by the Shenzhen Institute of Advanced Technology, Chinese Academy of Sciences (SIAT-IRB-200315-HO438).

**Large-scale cultivation of human stool-derived *in vitro* communities**

20ul stool slurries aliquot stocks were inoculated into 980 μL medium containing antibiotics in triplicate into 96 deep-well plates (PCR-96-SG-C, Axygen) for static culturing at 37 °C for 24h in the anaerobic workstation. The concentration for each antibiotic was evaluated as described in the SI method. The medium (MiPro) used for *in vitro* culture was modified from previous studies, which comprises: peptone water (2.0 g /L, CM0009, Thermo Fisher), yeast extract (2.0 g /L, LP0021B, Thermo Fisher), L-cysteine hydrochloride (1 g/ L), Tween 80 (2 mL/L), hemin (5 mg/L), vitamin K1(10 μL/L), NaCl (1.0 g /L), $K_2HPO_4$ (0.4 g/L), $KH_2PO_4$ (0.4 g/L), $MgSO_4 \cdot 7H_2O$ (0.1 g/L), $CaCl_2 \cdot 2H_2O$ (0.1 g/L), $NaHCO_3$ (4 g/L), porcine gastric mucin (4 g/L, M2378, Sigma-Aldrich), sodium cholate (0.25 g/L) and sodium chenodeoxycholate (0.25 g/L) (*6*). After 24h of antibiotics treatment, *in vitro* microbial communities were passaged every 24 h with a 1:200 dilution into fresh medium using the automated 96-format Thermo Scientific™ ClipTip™ (Thermofisher) pipette (every 24h, 5 μL of this saturated culture was transferred into 995 μL of fresh medium). After 5 days of passaging, 500 μL of the cultures were mixed with 500 μL sterile 40% glycerol (v/v, in sterile phosphate-buffered saline, with 0.1% L-cysteine hydrochloride) in crimp vials, sealed, and stored as baseline communities at −80 °C for further usage and long-term storage. After each transfer, the remaining samples were centrifuged to remove the supernatant, and the pellets were stored at -80°C with a plastic seal until DNA extraction. The *in vitro* microbial community biomass was evaluated by measurement of optical density ($OD_{600}$) with an Epoch 2 plate reader (BioTek) after 24h of incubation.

**Generation of baseline communities with diverse taxonomic profiles**

To examine if *in-vitro* stool-derived communities can reach stable states and display diverse compositions, we collected stool samples from healthy donors and grew them in MiPro medium, which has shown its capability in capturing and maintaining the diversity of *in vitro* stool-derived communities (*6-8*). We inoculated the stool aliquots into 96-well plates with growth media and incubated them in an anaerobic workstation in triplicate, passing them every 24h with a 1:200 dilution. The microbial communities were assessed by shallow metagenomic sequencing, which is a cost-effective method for characterizing species-level composition of microbiota samples (*9*). We collected time-series data to examine the dynamics of community establishment on the in-vitro platform. The metagenomic analysis revealed that, after an initial period of approximately four days, the composition profiles of almost all *in vitro* communities reached a stable and reproducible steady state. Our analysis also showed that the stool-derived *in vitro* communities were highly complex in their compositions and could retain personalized gut microbiota variation, as evidenced by species-level time-series compositions of 4 representative communities derived from 4 donors over ten rounds of *in vitro* passaging in MiPro (**Fig.S4A,B**).

From the fecal samples of SIAT cohort, we selected 24 donors in which *E. faecium and A. muciniphila* were not detected by metagenomic sequencing. To increase the diversity in baseline communities, we treated each donor's sample with 12 antibiotics from different classes (*10*) (**Fig.S2**). Different antibiotic classes target distinct spectra of bacteria, leading to a remodeling of the community in different directions (*10*). We selected antibiotics from different classes as described in the EUCAST databases (*11*). The optimal concentrations of the antibiotics were determined based on a previous study that evaluated the activity spectrum of antibiotic classes on human gut commensals (*10*). We tested at least three different concentrations for each antibiotic and evaluated the optimized dose based on its ability to partially inhibit (50%-80%) the overall growth of stool-derived bacteria as measured by $OD_{600}$ after 24h of incubation. To ensure reproducibility, we screened at least three different stool aliquot stocks as biological duplicates for each antibiotic. We measured the $OD_{600}$ of each well every 30 minutes using an Epoch 2 plate reader (BioTek) and collected growth curves up to 24h.

**Bacterial strains**

*Enterococcus faecium*, *Enterococcus faecalis* and *Clostridium symbiosum, Streptococcus* *salivarius* and *Bifidobacterium breve* strains were isolated from fecal samples of SIAT cohort. Taxonomy of isolates from SIAT cohort was confirmed by whole genome sequencing. Genome sequences have been deposited in PRJEB60398 (see data availability). *Lactobacillus plantarum HNU082 (12), Lactobacillus paracasei HNU312* (*13*) was provided by Prof. Jiachao Zhang from Hainan University. *Akkermansia muciniphila* (ATCC BAA-835) and *Fusobacterium nucleatum* (ATCC 25586) were purchased from ATCC.

**Profiling the colonization outcomes of different exogenous species**

We conducted a preliminary experiment to investigate the colonization outcome of gut microbial communities to different exogenous species (**Fig.S5**), including: *E. faecium, A. muciniphila* (*14*), *F. nucleatum, S. salivarius, B. breve* and *Lactobacillus spp.* (*L. plantarum HNU082* and *L. paracasei HNU312*). We identified 12 stool samples from healthy donors in which the selected invader species were undetectable in the microbiota. We then cultured the stool samples *in vitro* and exposed them to antibiotics before introducing the exogenous species (~5% of total biomass, approximately $10^6$ CFUs for each well) into the community. We used shallow metagenomic sequencing to monitor the time-series and final community composition.

**Invasion experiments of *E. faecium* and *A. muciniphila***

To conduct invasion experiments, frozen stocks of *E. faecium* (strain SIAT_DA797) and *A. muciniphila* (strain ATCC_BAA-835) were grown anaerobically in BHI and mGAM at 37℃, respectively, until stationary phase. *In vitro* microbial baseline communities, stored at −80°C, were thawed and revived by adding 20 μL of the stocks to 980 μL of MiPro medium in deep well plates. After incubation for 24 hours at 37°C, community biomass was measured by $OD_{600}$, and 5 μL of the saturated cultures were diluted into 1 mL of fresh MiPro in a new plate. Each well was invaded with the respective amount of *E. faecium* or *A. muciniphila*, with biomass representing 5% of the inoculated communities' average biomass. The inoculum was passaged every 24 hours of incubation, with a 1:200 dilution into fresh medium for 8-10 passages until the community reached a steady state (10 passages for *E. faecium,* 8 passages for *A. muciniphila,* based on data from Fig.S5). After each passage, the remaining samples were centrifuged to remove the supernatant, and the pellets were stored at -80°C with a plastic seal in plate until DNA extraction.

**Metagenomic sequencing and taxonomic profiling**

DNA was extracted from 200 mg of stool samples using the QIAamp Power Fecal Pro DNA Kit (Qiagen) according to the manufacturer's instructions. For stool-derived *in vitro*-cultured samples, 500 uL of cultured samples were used for DNA extraction with the DNeasy UltraClean 96 Microbial Kit (Qiagen) using an automated protocol at Tecan Freedom EVO 200. The Hieff NGS® OnePot II DNA Library Prep Kit for Illumina® (Yeasen) was used for library preparation, following the manufacturer's instructions. The resulting library DNA was cleaned up and size-selected with Hieff NGS® DNA Selection Beads (Yeasen), and quantified using the dsDNA High Sensitivity kit on a Qubit (Thermo Fisher). Libraries were further pooled together at equal molar ratios, and the purity and library length distribution were assessed using Bioanalyzer High Sensitivity DNA Kit (Agilent). Sequencing was performed on the Illumina HiSeq X Ten system (150bp paired-end reads; Annoroad Gene Technology Co.), with a target sequencing depth of 0.3 Gbp raw data per sample, as recommended by previous studies (*9*).

Samples with fewer than $10^5$ clean reads were excluded from downstream analysis. Prior to analysis, reads were trimmed using the following criteria: (1) Removing reads with more than 50% of the base below quality score 19; (2) Removing reads with more than 5% of the base being N; (3) Discarding paired-end reads if either of the paired reads did not meet the above criteria. Microbial community composition from metagenomic sequencing data was generated using the SHOGUN pipeline and the RefSeq database version 82, as described in previous studies (*9, 15*). Species-level abundance profiles were filtered by using a relative abundance threshold of 0.0001 (0.001) for all taxa in colonization prediction of *E. faecium* (*A. muciniphila*), and those low-prevalence taxa (present in less than 20% samples) were further filtered to reduce the feature number. The colonization outcomes were evaluated based on the invader's absolute abundance in the community, which was estimated by multiplying the relative abundance and the $OD_{600}$ value ($OD_{600}$×relative abundance). To ensure repeatability, samples with Pearson correlation below 0.8 among replicates were excluded from COP analysis. This resulted in the exclusion of 1.8% of samples for *E. faecium* and 1.3% for *A. muciniphila.*

**Quantification of the relative abundance of *E.faecium* and *A.muciniphila* by metagenomic sequencing**

To confirm the accuracy of shallow metagenomic sequencing in quantifying the relative abundance of *E. faecium* and *A. muciniphila*, a spike-in experiment was conducted (**Fig.S18A**). In this experiment, a predefined amount of bacterial DNA from the target species was added to a metaDNA sample extracted from an *in vitro* community derived from human stool. This metaDNA sample was used as the background, since it has been previously sequenced and did not contain the target species. The spike-in DNA of the target species (*E. faecium* or *A. muciniphila*) was 1:10 diluted for eight times and was added to the microbial metaDNA to a mixed DNA sample (5 μL of target species DNA into 30 ng of microbial metaDNA). Three replicates were made for each sample. The mixed DNA was then used for library construction and metagenomic sequencing. By comparing the detected relative abundance generated by shallow metagenomic sequencing with the expected abundance, the accuracy and sensitivity of our workflow were determined. The detection threshold of *E. faecium* is 0.0001 (**Fig.S18B**) and the detection threshold of *A.*

*muciniphila* is 0.001 (**Fig.S18C**). Our results showed that the quantification of the relative abundance of the two target species using the shallow metagenomic sequencing pipeline is accurate and reproducible.

**Colonization impact of resident species onto the invading species**

To compute the colonization impact, e.g., the impact of resident species onto the colonization outcome of the invading species, we first trained the prediction models using all the samples. Then, for resident species $i$ in a permissive local community $\alpha$, we performed a thought experiment by introducing a perturbation in the abundance of resident species $i$, and used the trained machine learning model to predict the new steady state abundance of invading species $\tilde{x}_i^\alpha$ after the perturbation. The colonization impact (CI) of resident species $i$ onto the invading species in local community $\alpha$ is defined as:

$$\mathrm{CI}_i^\alpha = \frac{\tilde{x}_i^\alpha - x_i^\alpha}{\tilde{x}_i^\alpha + x_i^\alpha}$$

where $x_i^\alpha$ is the steady state abundance of invading species in community $\alpha$ before perturbing the abundance of species $i$. A negative colonization impact ($\mathrm{CI}_i^\alpha < 0$) indicates that species $i$ inhibits the colonization of the invading species in community $\alpha$. For classification models, $x_i^\alpha$ and $\tilde{x}_i^\alpha$ represent the colonization probability before and after perturbing the abundance of species $i$.

**Validation of the inhibitory effect of *E. faecalis* on *E. faecium* colonization**

<u>Pairwise co-culture experiments</u>

Soft Agar Overlay Assays were conducted using BHI agar plate. *E. faecium* DA797 was cultured to an $OD_{600}$ of 0.6 and 100ul of the inoculum was pipetted into 10mL prewarmed (42°C) BHI containing 0.75% (w/v) agar. The mixture was briefly mixed and then transferred onto a plate already laid with 10mL BHI 1.5% agar and four Oxford cups, to embed *E. faecium* into soft agar. The mixture was spread evenly on the surface of the plate. Next, 100-µl volumes of *E. faecium, E. faecalis* DA894, *E. faecalis* DA462 ($OD_{600}$=0.6) were added individually into the Oxford cups. The plates were incubated anaerobically at 37 °C for 24h before observation. The experiment was performed three times with two technical replicates for each strain.

Liquid co-culture experiments were performed in BHI at 37°C static, under anaerobic conditions. *E. faecium* and *E. faecalis* were cultured separately in BHI at 37°C for 24h without shaking, then diluted in BHI to an $OD_{600}$ of 0.005 and then inoculated at 1:1 ratio into 1 mL of BHI broth and grown for 24h without shaking. Mono- and co-culture outputs were centrifuged to remove the supernatant, and the pellets were subsequently DNA extracted and *E. faecium* specific qPCR primer was used to detect the abundance of *E. faecium*.

Community experiments

Frozen stocks of *E. faecium* DA797, *E. faecalis* DA462 and DA894 and *C. symbiosum* DA229, were grown anaerobically at 37 °C in BHI until they reached the stationary phase. Eight baseline communities' stocks were revived into 980µL MiPro medium with three replicates in deep well plates. After 24h's incubation at 37 °C, the community biomass was measured by $OD_{600}$. Saturated cultures were then diluted 5µL into 1mL of fresh MiPro in a new 96-well plate before the invasion experiments. Three different experimental schemes were used: 1) Add *E. faecalis* (or *C. symbiosum*) into the baseline community, followed by *E. faecium* on the next day; 2) Add *E. faecalis* and *E. faecium* on the same day; 3) Add *E. faecium* into the baseline community, followed by *E. faecalis* on the next day. The inoculum was incubated at 37 °C and serially diluted every 24 h of 7 passages until the community reached a steady state. Saturated cultures were centrifuged to remove the supernatant, and the pellets were stored at -80°C with a plastic seal until DNA extraction. *E. faecium* abundance was assessed by both metagenomic sequencing and qPCR.

qPCR assays for absolute quantification

qPCR reactions were used to validate the impact of *E. faecalis* on the colonization outcome of *E. faecium*. qPCR reactions (0.5 µl DNA, 0.2 µM each primer, Hieff® qPCR SYBR Green Master Mix (Yeasen) were performed on a Bio-Rad CFX384 Touch Real-Time PCR Detection System, using primers specific for *E. faecium* under the following reaction conditions: 95 °C for 5min followed by 40 cycles of 95 °C for 10s , 60°C for 20 s and 72°C 20 s. *E. faecium*-specific primer sequences were: Ala-F:ATCCCTCTGGGCACGCAC, Ala-R:ACATACACGCCCAATCGTTTC, as described previously (*16*). Standard curves using genomic DNA of *E. faecium* were used for absolute quantification of *E. faecium* copy numbers.

#### *E. faecium* and *E. faecalis* abundance analysis in human cohorts

The following datasets were used for the metagenomic analysis of the species of interest in four large and diverse human cohorts: Israel (*17*), Lifelines-DEEP (*18*), PERDICT-1 (*19*), TwinsUK (*20*) and SIAT cohort. Sequencing data were obtained using the accession numbers provided in the associated references and processed by SHOGUN pipeline as previously described. *E. faecium* and *E. faecalis* with relative abundance below 0.0001 is set to $10^{-4}$ for visualization.

#### Statistical analysis

Statistical details for each experiment are indicated in the figure legends. Pearson correlation coefficients and the p-values for testing replicates communities' composition correlation were calculated on $\log_{10}$(relative abundance). Kendall correlation coefficients and the p-values for testing *E. faecium* and *E. faecalis* abundance correlation were calculated on $\log_{10}$(relative abundance). Alpha diversity of the community was calculated on species profile using the observed species richness and Shannon index. The composition of microbiota and variations in colonization outcomes between communities were analyzed by performing PCoA using the Bray-Curtis dissimilarity metric on the species-level abundance profile. Similarities among groups were determined by permutational multivariate analysis of variance (PERMANOVA, Adonis test) based on the Bray-Curtis dissimilarity (*21*), with 999 permutations used to test the significance. These analyses were conducted using the vegan (*22*) package (version 2.6-4). Non-parametric Mann-Whitney U-test were used to conduct pairwise comparisons between two groups (*23*). P values of less than 0.05 were considered as statistically significant, as indicated in the figures (ns, not significant, *p<0.05, **p<0.01, ***p<0.001, ****p<0.0001). Data analysis and plotting was performed in R version 4.1.2 and R studio version 2022.12.0+353 using the packages dplyr, ggpubr, vegen and ComplexHeatmap.

### Supplementary Text

**Analytical derivation on the steady state abundance of exogenous species in GLV model**

For a local community $\alpha$ of $s$ resident species governed by GLV dynamics, we denote the post-invasion steady state abundance of the exogenous species as $x_{s+1}^{(1)}$. After invasion, the community arrives at a new steady state, i.e., $\frac{dx_{s+1}(t)}{dt} = 0$. Thus, according to Eq.1, $x_{s+1}^{(1)}$ can be expressed as:

$$x_{s+1}^{(1)} = r_{s+1} + \boldsymbol{c}\mathbf{x}_{1:S}^{(1)} \quad (2)$$

Here, the $s$-dimensional vector $\boldsymbol{c}$ represents the interaction strength of the resident species onto the exogenous species, $\mathbf{x}_{1:S}^{(1)}$ represents the post-invasion steady state abundance of the resident species.

Based on derivations in our previous study (*24*), the shift in the steady state abundance of resident species (i.e. the difference between $\mathbf{x}_{1:S}^{(1)}$ and the pre-invasion steady state $\mathbf{x}_{1:S}^{(0)}$) satisfies the following relation:

$$\mathbf{x}_{1:S}^{(1)} - \mathbf{x}_{1:S}^{(0)} = -\boldsymbol{A}^{-1}\boldsymbol{b}x_{s+1}^{(1)} \quad (3)$$

Here, the $s$-dimensional state vector $\mathbf{x}_{1:S}^{(0)} = [\mathbf{x}_1^{(0)}, \mathbf{x}_2^{(0)}, \cdots \mathbf{x}_S^{(0)}]^{\mathrm{T}}$ represents the pre-invasion steady state of the local community, the $s$-dimensional vector $\boldsymbol{b}$ represents the interaction strength of the exogenous species onto the resident species. The interactions among resident species are encoded in matrix $\boldsymbol{A}$.

By combining Eq.2 and Eq.3, we discovered a surprisingly simple linear relation between the post-invasion abundance of the exogenous species $x_{s+1}^{(1)}$ and the pre-invasion abundance of resident species $\mathbf{x}_{1:S}^{(0)}$:

$$x_{s+1}^{(1)} = \frac{r_1 + \boldsymbol{c}^{\mathrm{T}}\mathbf{x}_{1:S}^{(0)}}{1 + \boldsymbol{c}^{\mathrm{T}}\boldsymbol{A}^{-1}\boldsymbol{b}} \quad (4)$$

The analytically derived relation can fully explain the simulated colonization outcomes in GLV model (**Fig.S1**, Spearman correlation $\rho = 1, p < 0.001$). Although the linear relation in Eq. 4 doesn't hold for other dynamical models (e.g., non-linear interactions), it gives us important insights that learning the mapping for colonization outcome prediction is feasible by data-driven models and the number of parameters required for fitting the relation is on the order of *~O(N)*. This is consistent with our observations on the number of training samples required for accurate prediction of colonization outcomes (**Fig. 1**).

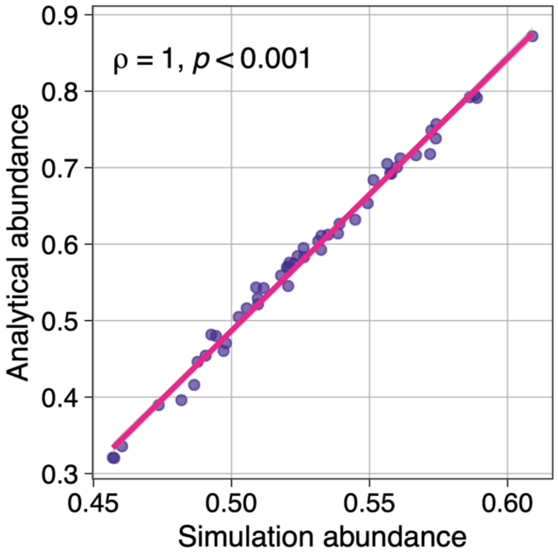


**Fig. S1. The steady state abundance of an invading species in communities governed by GLV dynamics: comparison between analytical derivations and simulations.** The analytically derived relation (Equation 4 in **Supplementary Text**) can fully explain the simulated colonization outcomes in GLV model (Spearman correlation $\rho = 1, p < 0.001$). We generated 50 local communities, each consisting of 4 species randomly drawn from a meta-community of 7 species. Network connectivity $C = 1$ and interaction strength $\sigma = 0.2$. Species-8 was introduced as an invading species.

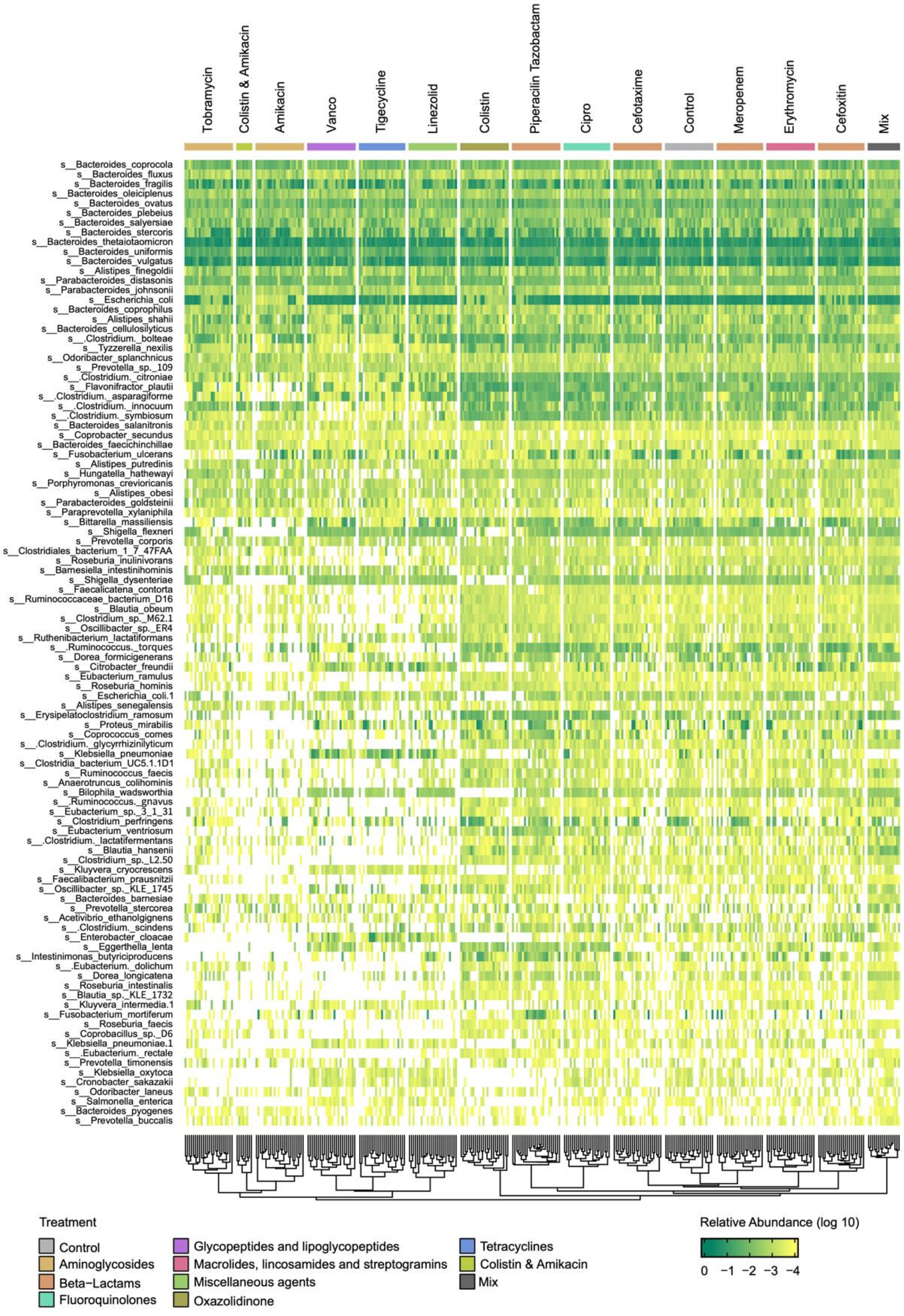


**Figure S2. The compositional profile of baseline communities at the species level.** Each column corresponds to a baseline community derived from a human stool sample (24 donors) treated with antibiotics (12 antibiotics). Mix indicates the group of communities derived from mixing two different donors. Each row corresponds to a species, clustered by the similarity of relative abundance across baseline communities. Species with top 100 prevalence are displayed.

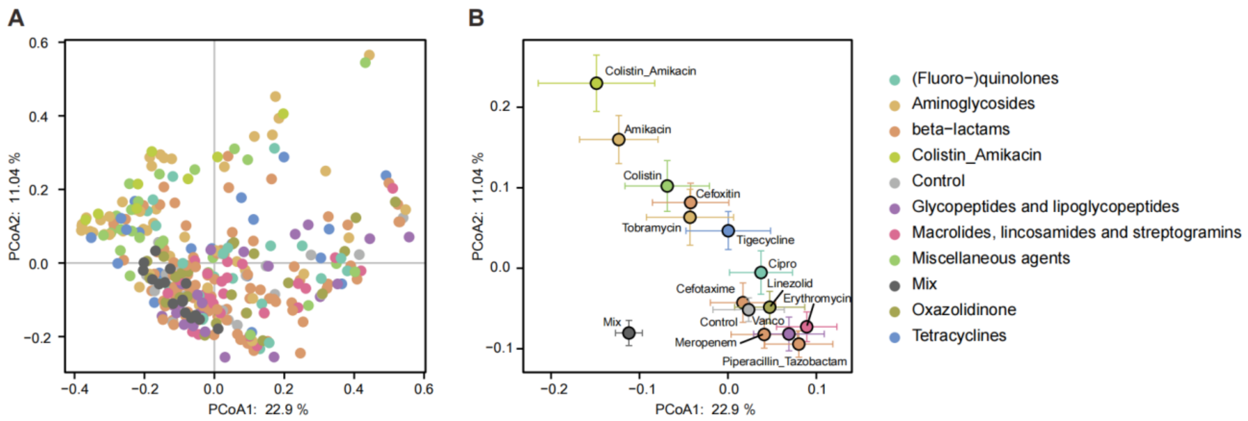


**Figure S3. Generation of diverse baseline communities by antibiotics treatments. (A)** Principal-coordinate analysis (PCoA) based on the Bray-Curtis dissimilarity of the compositional profiles at the species level. The baseline communities are color-coded according to antibiotics treatments. **(B)** The colored dot for each antibiotics treatment represents the compositional profile averaged over 24 subjects. Error bars are SEMs. The antibiotics of different classes had distinct impacts on community structure. Tobramycin and amikacin, belonging to aminoglycosides, drastically changed the community structure. In contrast, meropenem, cefoxitin, and cefotaxime, belonging to beta-lactams, had relatively moderate impacts on the community structure.

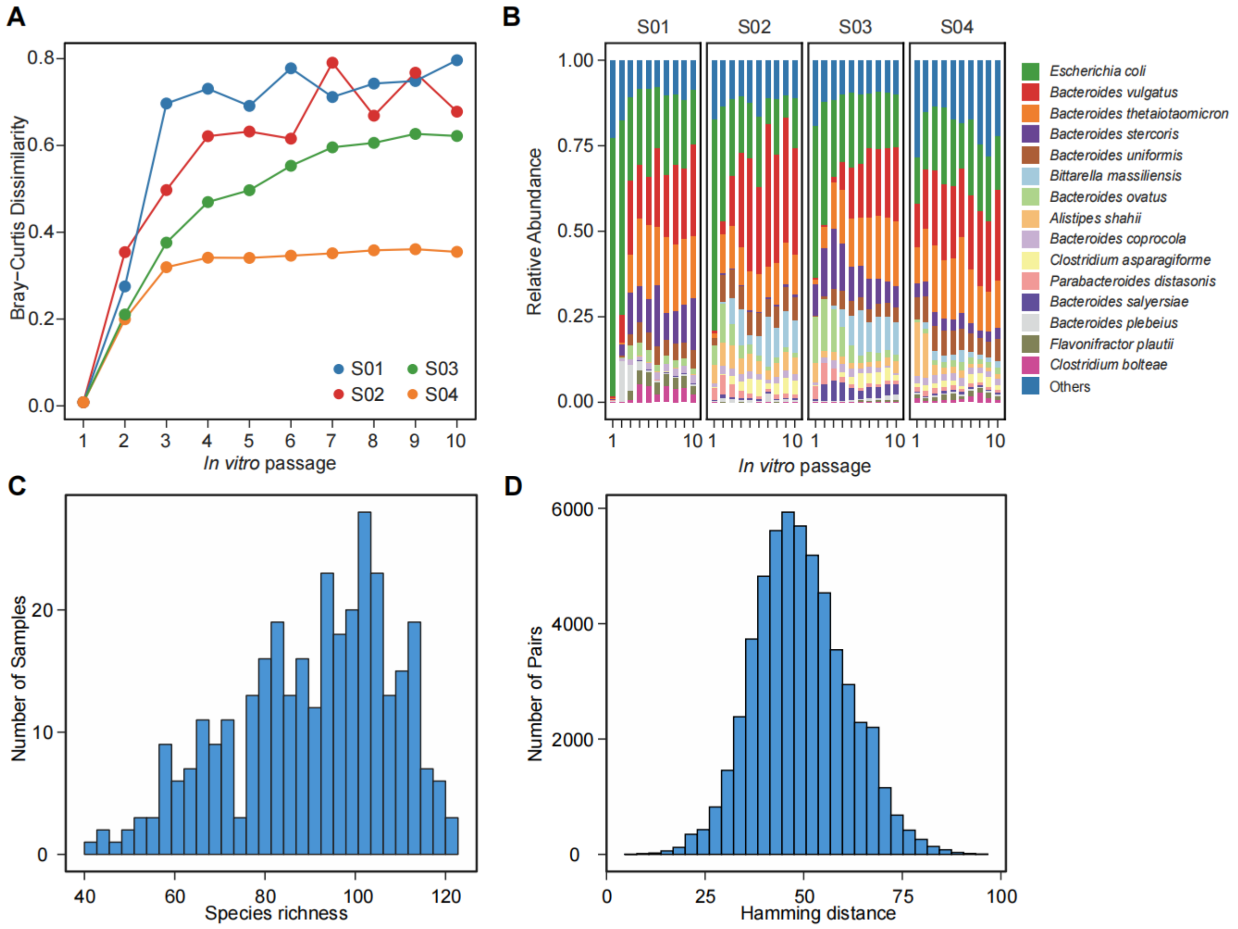


**Figure S4. Stabilization of human stool-derived *in vitro* communities and the statistics of steady-state baseline community composition. (A)** The Bray-Curtis dissimilarity to the initial compositional profile during serial passaging. Colored lines indicate the trajectories of communities from different donors (S01-S04). **(B)** Time series of the compositional profiles. The human stool-derived *in vitro* communities reached steady states after ~5 rounds of serial passaging in the MiPro medium. **(C)** Species richness of steady-state baseline communities. **(D)** Hamming distance between the species presence/absence profiles of baseline communities.

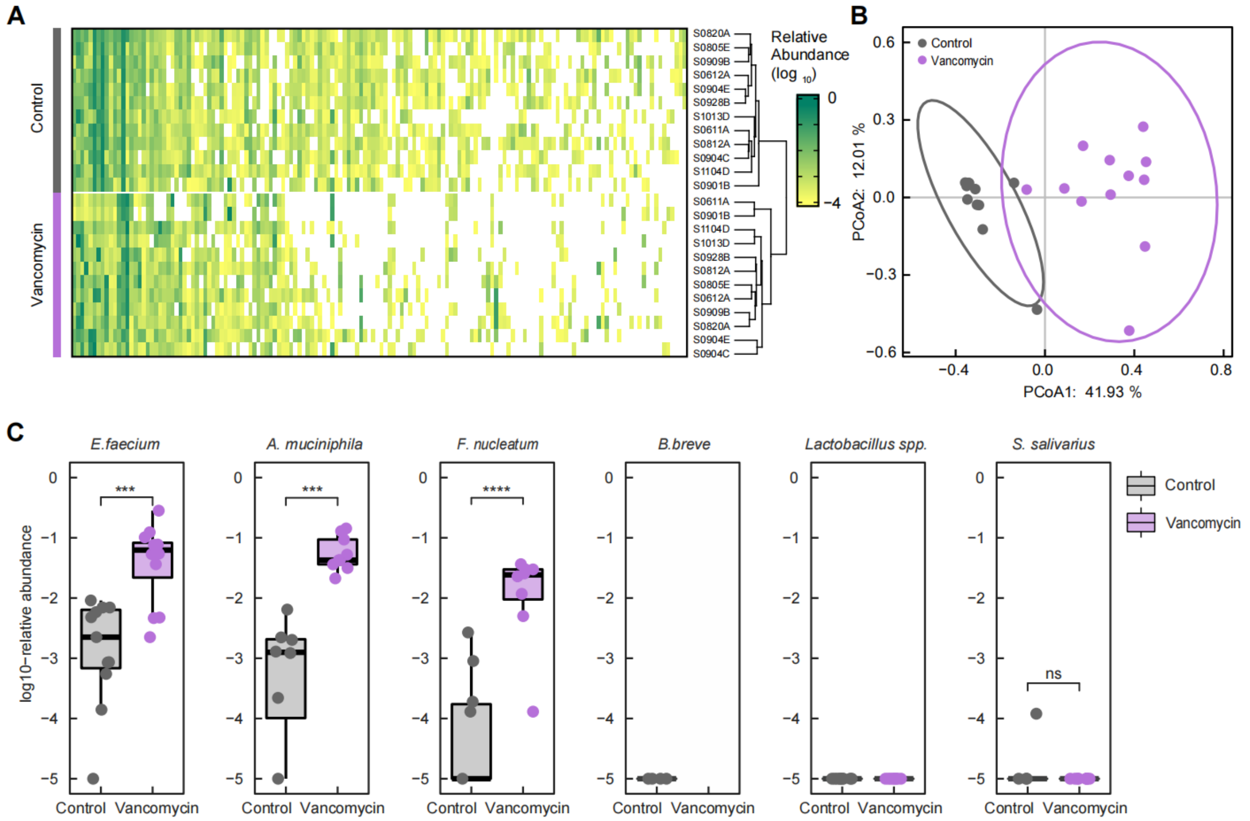


**Fig. S5. Colonization outcomes of different exogenous microbial species in human stool-derived *in vitro* communities. (A)** The compositional profile of baseline communities at the species level. Each row corresponds to a baseline community derived from a human stool sample (12 donors) treated with vancomycin (Vanco) or not (Control). Each column corresponds to a species, clustered by the similarity of relative abundance across baseline communities. Species with top 100 prevalence are displayed. **(B)** Vancomycin treatment altered the community structure at the species level (Adonis test, $R^2$=0.36，p<0.0001 ), as determined by PERMANOVA based on the Bray-Curtis dissimilarity. **(C)** Colonization outcomes of different exogenous species, including *E. faecium, A. muciniphila, F. nucleatum, S. salivarius, B. breve* and *Lactobacillus spp.* (*L. plantarum HNU082* and *L. paracasei HNU312*). The relative abundance of the invading species was determined by metagenomic sequencing of the final time point. We found that *E. faecium*, *A. muciniphila* and *F. nucleatum* could successfully colonize in some communities at varying levels of post-invasion abundance. In contrast, *S. salivarius, B. breve* and *Lactobacillus spp.* were unable to colonize in nearly all the baseline communities that we tested. Moreover, we found that vancomycin treatment significantly altered the colonization outcomes, rendering the communities more susceptible to invasion. ns, not significant, ***p < 0.001, ****p < 0.0001, Mann-Whitney U-tests. For visualization, the relative abundance was set to $10^{-5}$ if it was below the detection limit (i.e. failed invasion).

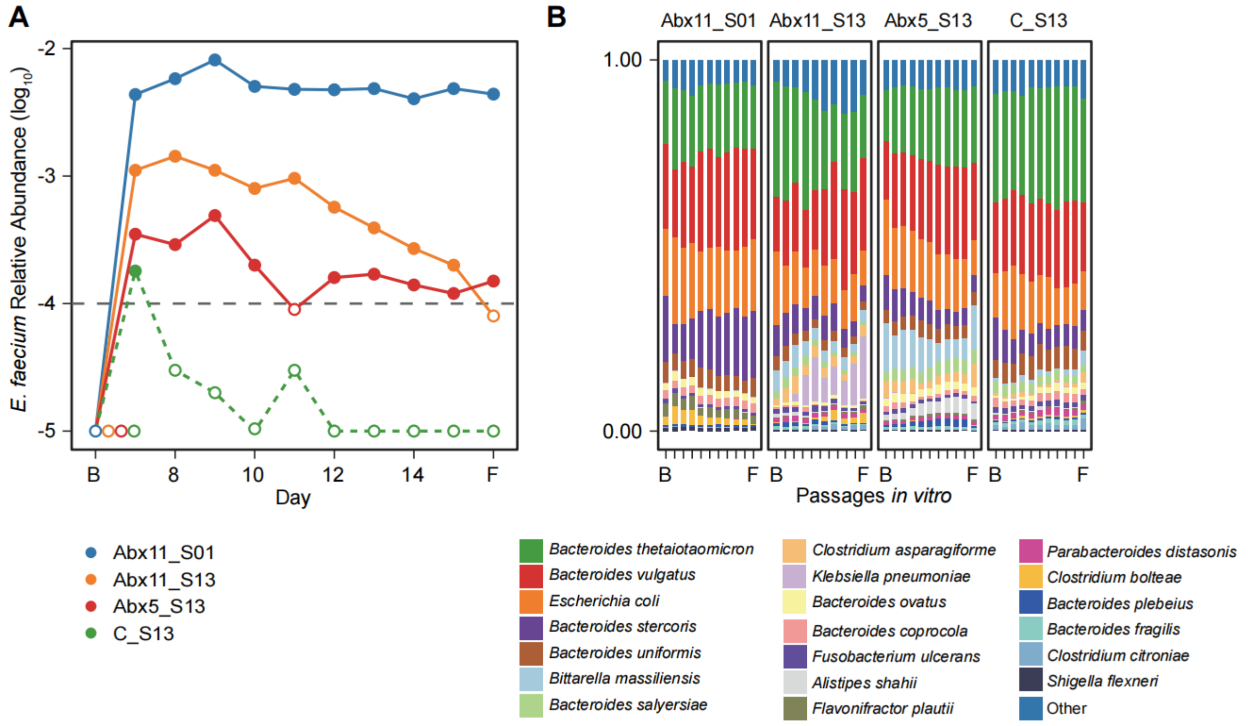


**Fig. S6. Post-invasion time series of *E. faecium* abundance and community composition. (A)** The colonization outcome of *E. faecium* in different communities was persistent during serial passaging. The dashed line indicates the detection limit of the relative abundance of *E. faecium* (Fig.S18). **(B)** The community composition was stable during serial passaging. B and F denote the baseline and the final time point.

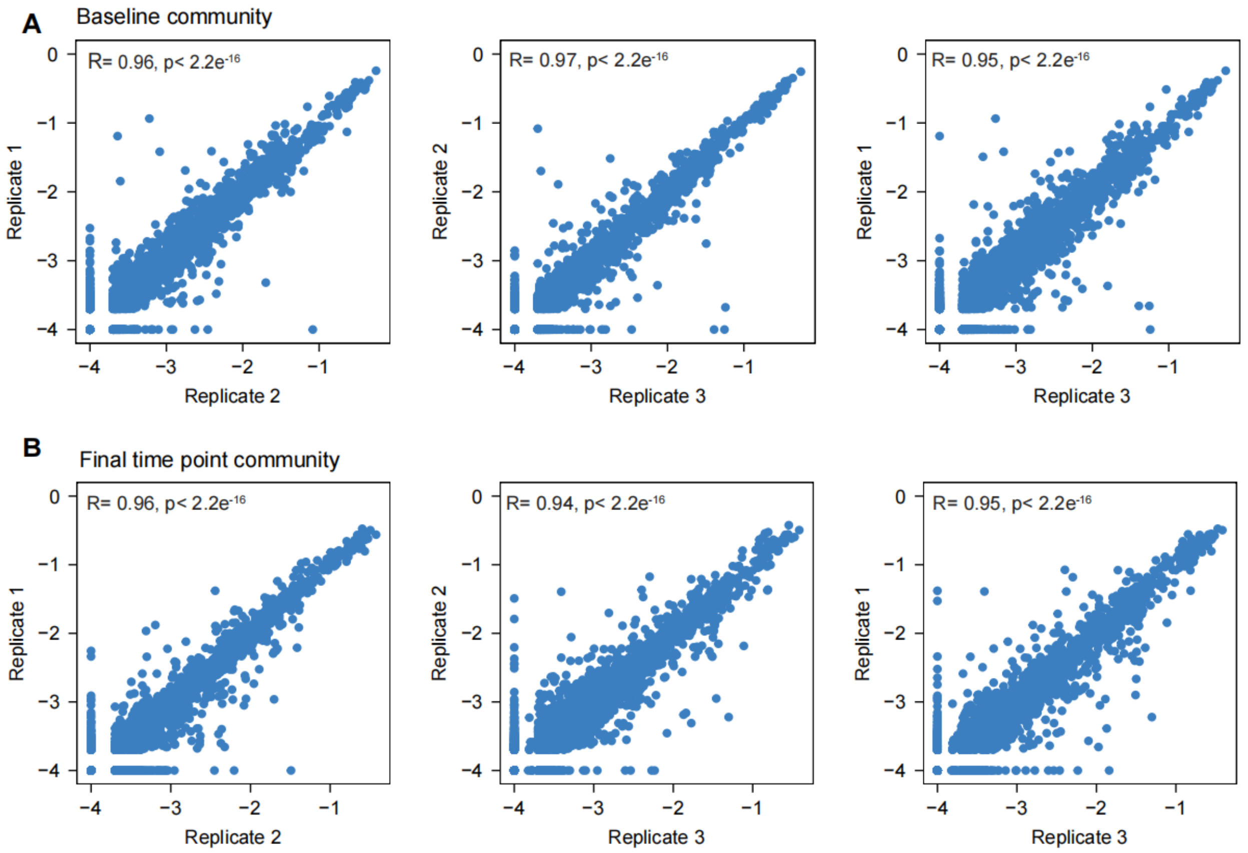


**Fig. S7. The composition of *in vitro* communities before and post *E. faecium* invasion is highly reproducible across replicates.** The species-level compositional profile of the baseline communities (**A**) and of the post-invasion communities (**B**) is highly reproducible among technical replicates (Pearson correlation). For visualization, the relative abundance was set to $10^{-4}$ if it was below the detection limit. n= 3 replicates.

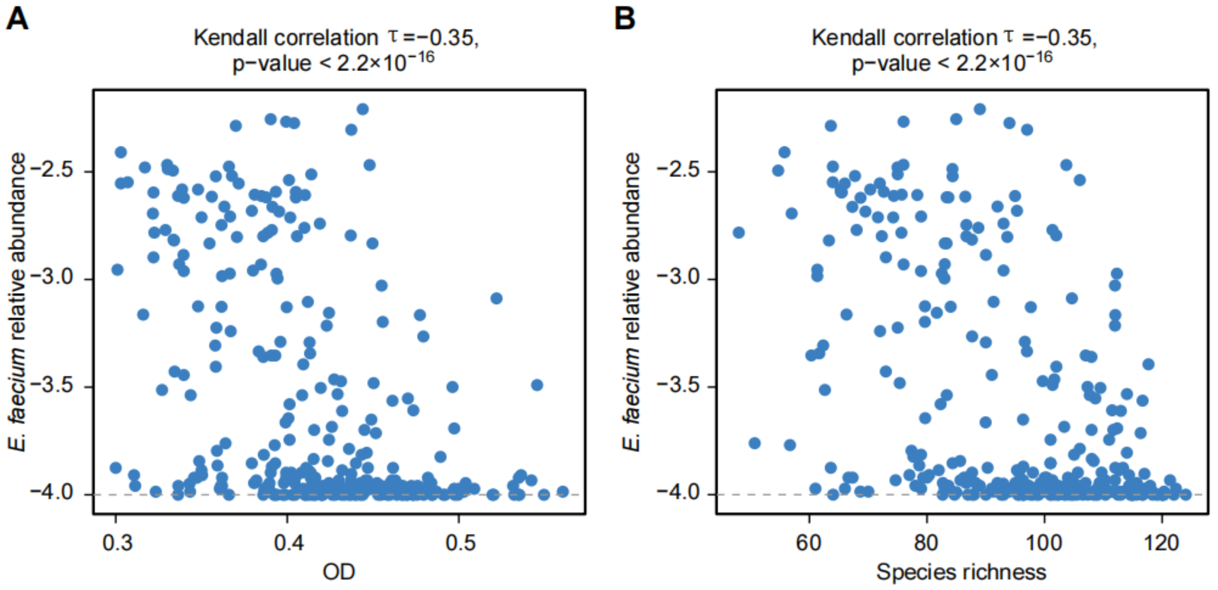


**Fig. S8. The invasion resistance to *E. faecium* increases with community biomass and diversity. (A)** The post-invasion steady state abundance of *E. faecium* is negatively correlated with the biomass of baseline communities (measured by $OD_{600}$). **(B)** The post-invasion steady state abundance of *E. faecium* is negatively correlated with the species richness of baseline communities.

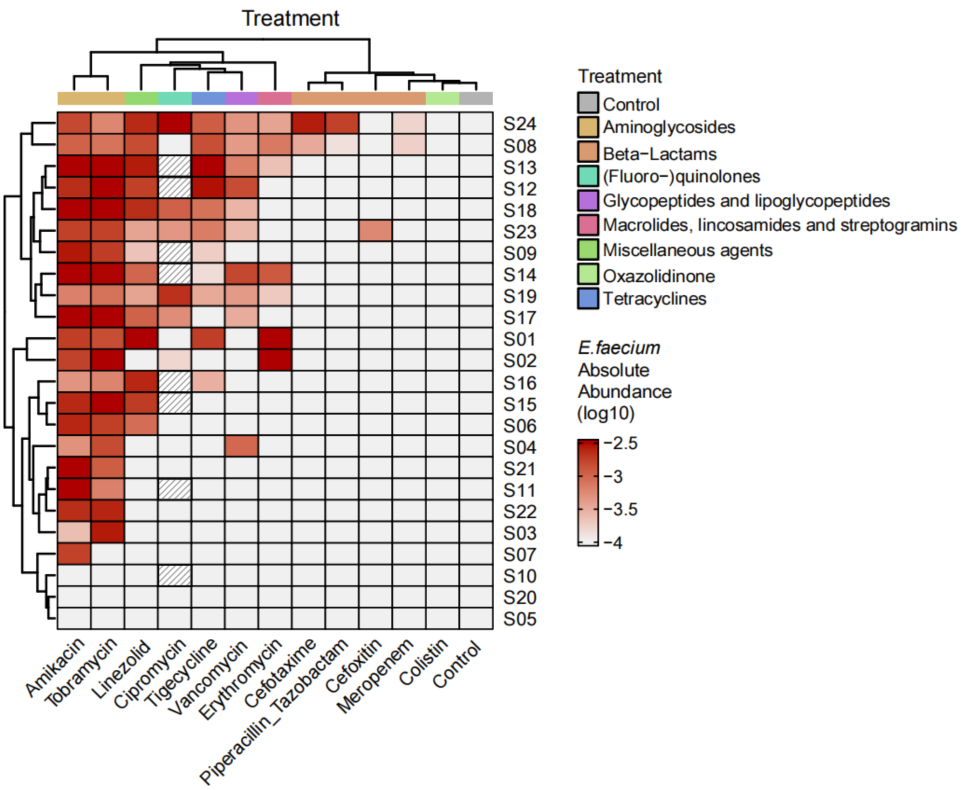


**Fig. S9. Variations in the colonization outcomes of *E. faecium* across different donors and antibiotics treatments.** For instance, post-invasion abundance of *E. faecium* in communities derived from donor S24 was higher than other donors; post-invasion abundance of *E. faecium* in communities treated with amikacin was higher than the control group and other treatment groups. Each row corresponds to a donor from which the communities were derived, each column corresponds to a treatment. The color gradient represents absolute abundance ($OD_{600}$×relative abundance) of *E. faecium* at the post-invasion steady state. Samples marked with slashes are not available.

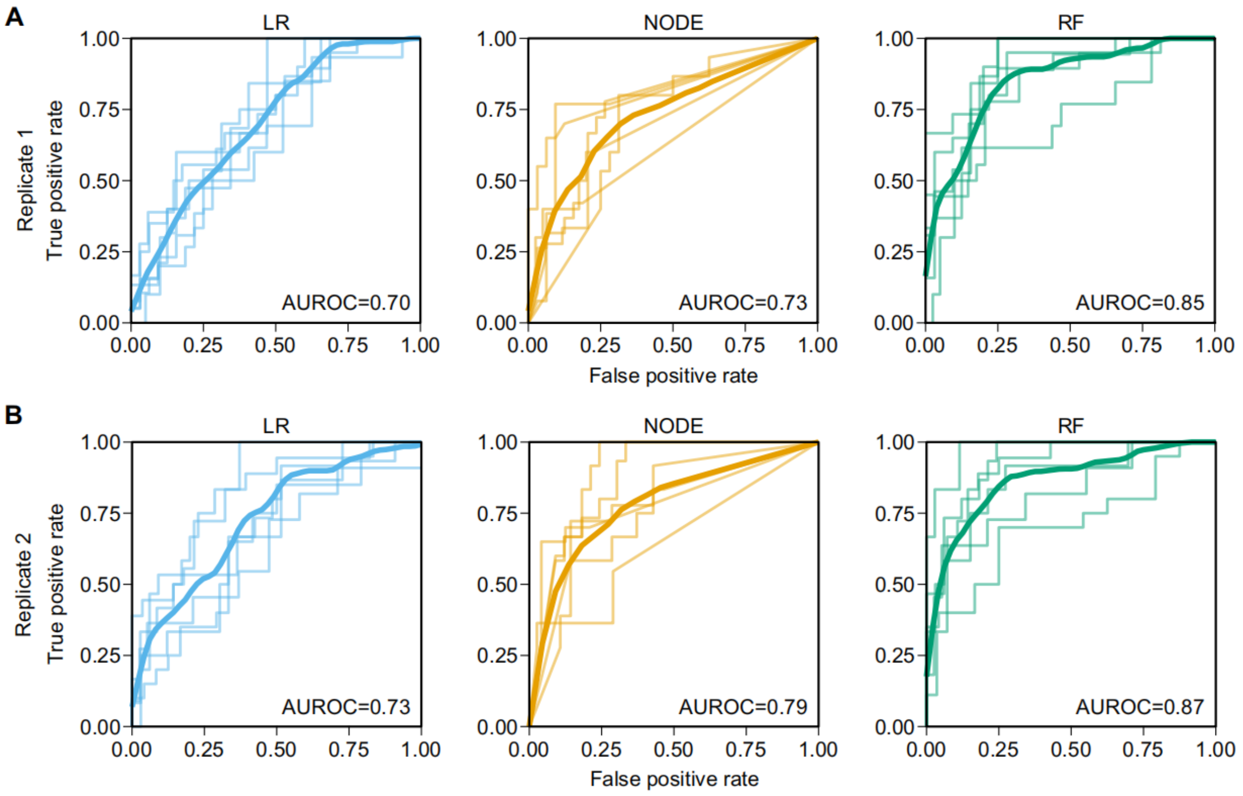


**Fig. S10. The performance of colonization outcome prediction for *E. faecium* is consistent across replicates.** ROC curve of machine learning models in binary classification (permissive vs. resistant) of the colonization outcomes of *E. faecium* in replicate 1 **(A)** and replicate 2 **(B)**. For each 6-fold cross validation (ROC curves shown in light color), we trained each model using the samples from 20 subjects and the samples from the remaining 4 subjects to evaluate the model. The mean ROC curve is shown in dark color. LR: Logistic Regression, NODE: COP-Neural Ordinary Differential Equations classifier, RF: Random Forest classifier.

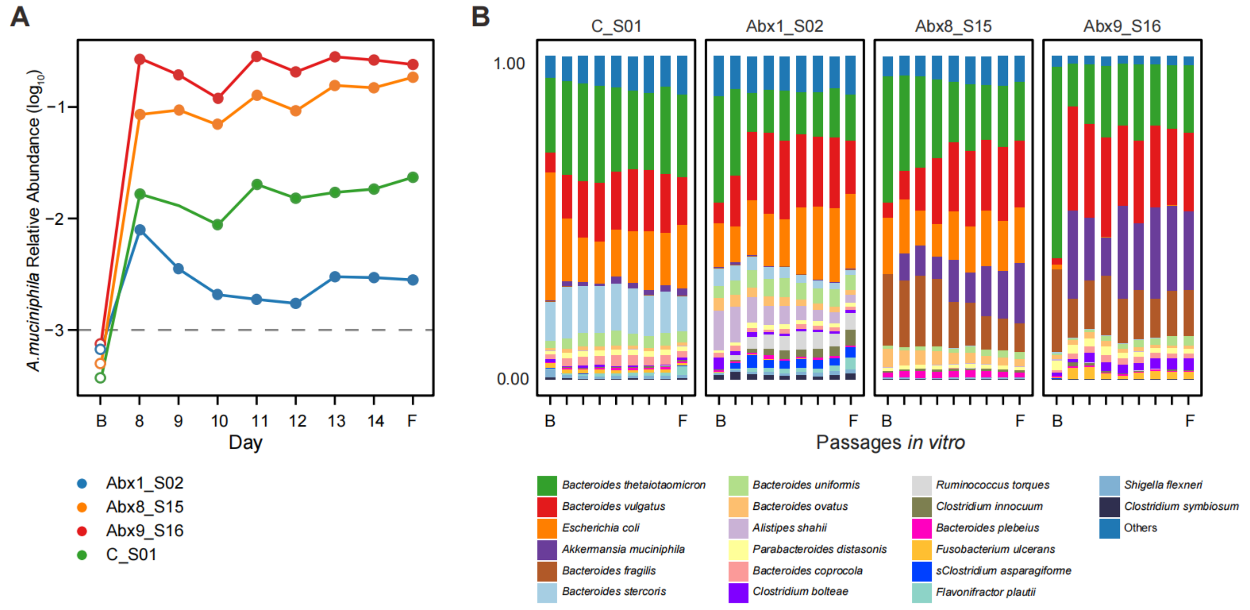


**Fig. S11. Post-invasion time series of *A. muciniphila* abundance and community composition.** **(A)** The colonization outcome of *A. muciniphila* in different communities was persistent during serial passaging. The dashed line indicates the detection limit of the relative abundance of *A. muciniphila* (Fig.S18). **(B)** The community composition was stable during serial passaging. B and F denote the baseline and the final time point.

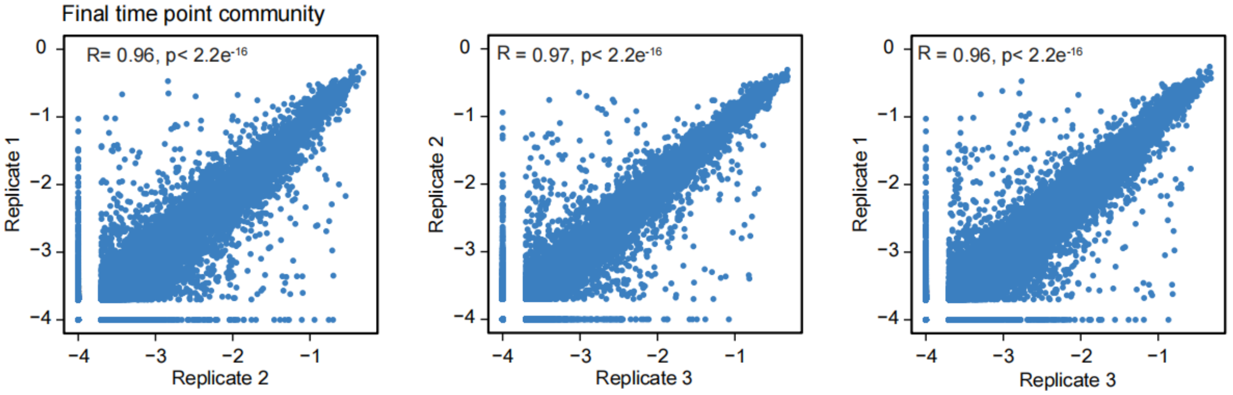


**Fig. S12. The composition of *in vitro* communities post *A. muciniphila* invasion is highly reproducible across replicates.** The species-level compositional profile of the post-invasion communities is highly reproducible among technical replicates (Pearson correlation). For visualization, the relative abundance was set to $10^{-4}$ if it was below the detection limit. n= 3 replicates.

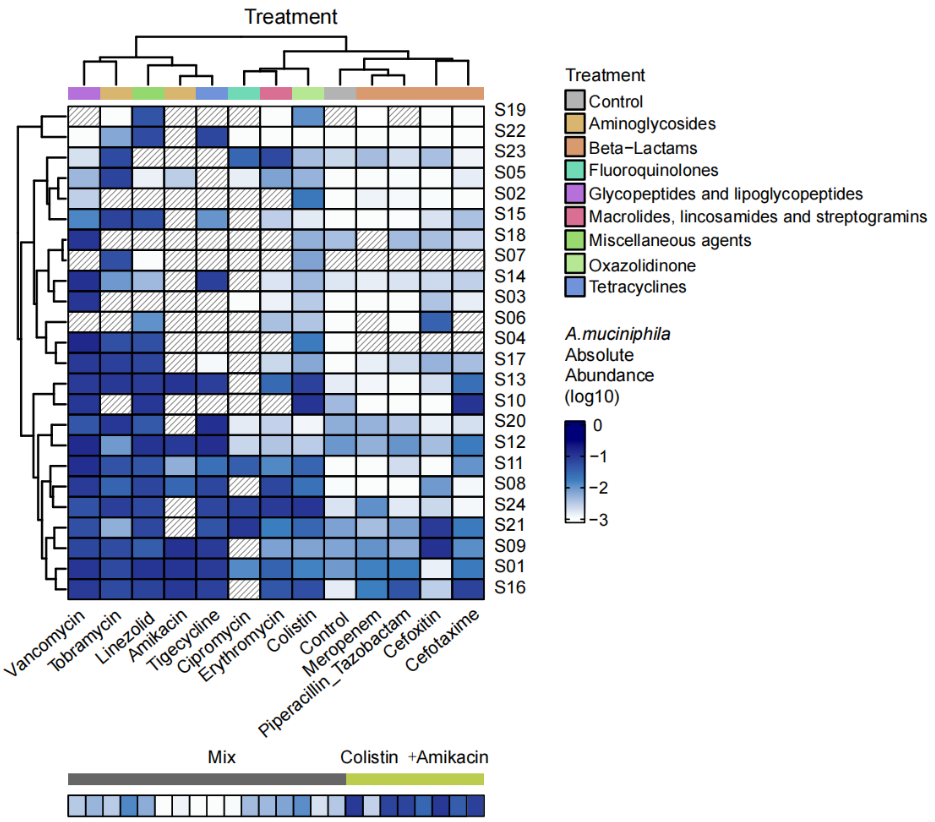


**Fig. S13. Variations in the colonization outcomes of *A. muciniphila* across different donors and antibiotics treatments.** For instance, post-invasion abundance of *A. muciniphila* in communities derived from donor S16 was higher than other donors; post-invasion abundance of *A. muciniphila* in communities treated with vancomycin was higher than the control group and other treatment groups. Each row corresponds to a donor from which the communities were derived, each column corresponds to a treatment. The color gradient represents absolute abundance ($OD_{600}$×relative abundance) of *A. muciniphila* at the post-invasion steady state. Samples marked with slashes were not used in *A. muciniphila* invasion experiments. Mix indicates the group of communities derived from mixing two different donors.

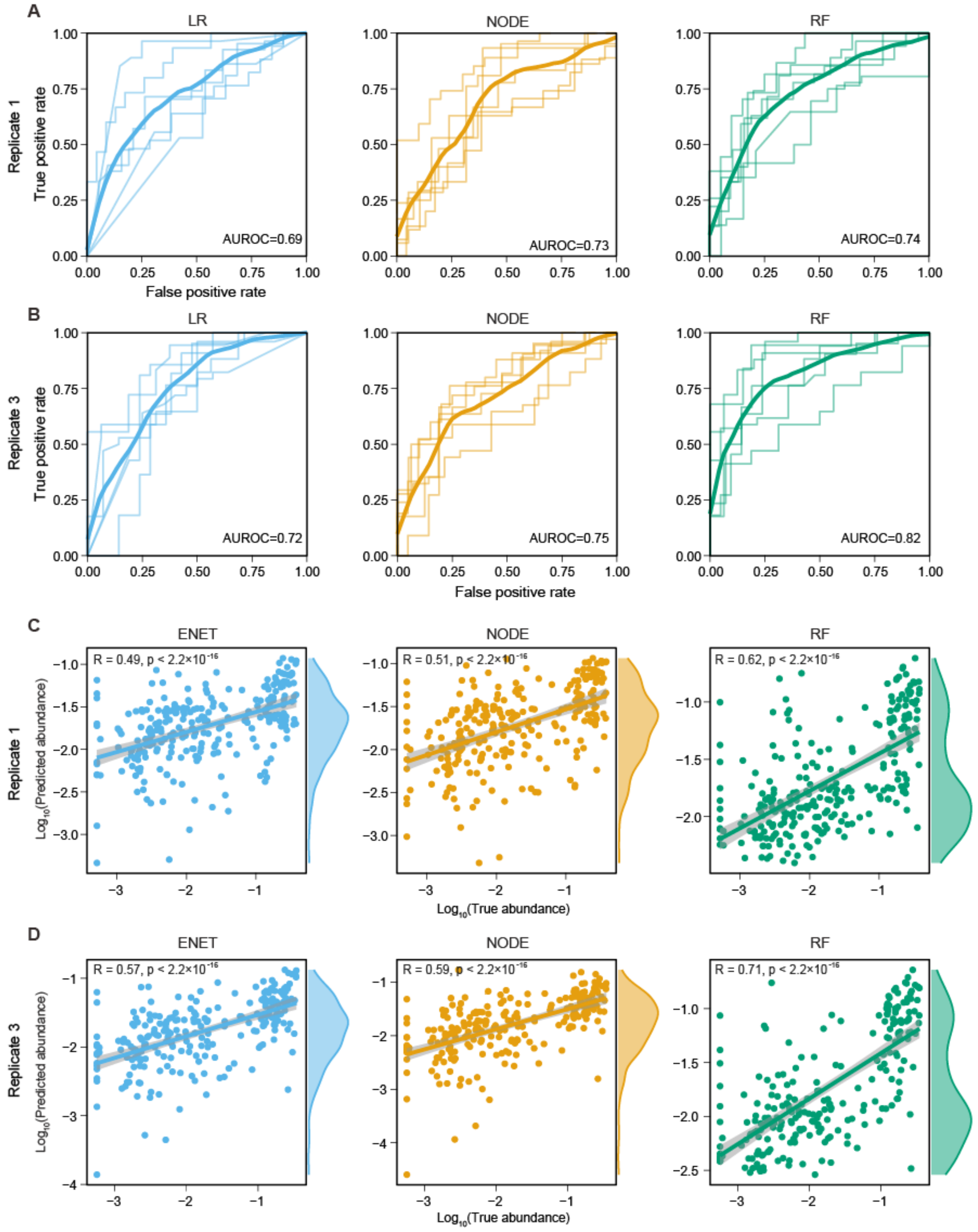


**Fig. S14. The performance of colonization outcome prediction for *A. muciniphila* is consistent across replicates.** ROC curve of machine learning models in binary classification (high permissive vs. Low permissive) of the colonization outcomes of *A. muciniphila* in replicate 1 **(A)** and replicate 3 **(B)**. For each 6-fold cross validation (ROC curves shown in light color), we trained each model using the samples from 20 subjects and the samples from the remaining 4 subjects to evaluate the model. The mean ROC curve is shown in dark color. Pearson's correlation coefficient between the predicted abundance and the true abundance of *A. muciniphila* in replicate 1 **(C)** and replicate 3 **(D)**. LR: Logistic Regression, ENET: Elastic Net Linear Regression, NODE: COP-Neural Ordinary Differential Equations regressor, RF: Random Forest regressor.

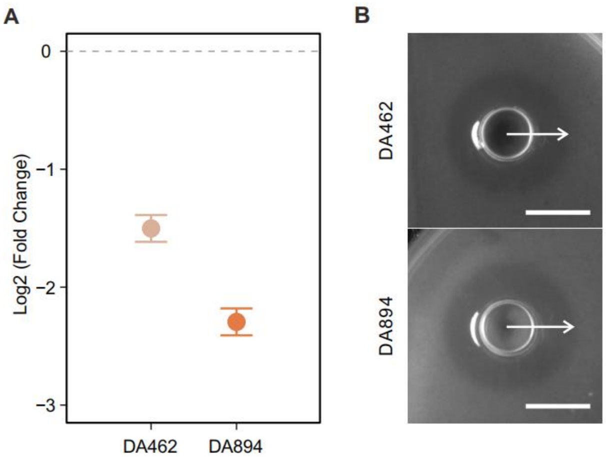


**Fig. S15. *E. faecalis* inhibits the growth of *E. faecium* in pairwise co-culture. (A)** The fold change in the abundance of *E. faecium* (the pairwise co-culture group divided by the mono-culture group) was lower than 1 (dashed line), indicating that the growth of *E. faecium* was inhibited in the presence of *E. faecalis* during pairwise co-culture in BHI. n= 3 replicates, the error bars are SEMs, measured by qPCR. **(B)** The Oxford cup assay was used to determine the inhibition of *E. faecium* by *E. faecalis.* An inhibition zone surrounding the Oxford Cup when *E. faecalis* was present. Scale bar, 1 cm. Two *E. faecalis* strains DA462 and DA894 were used in the assays.

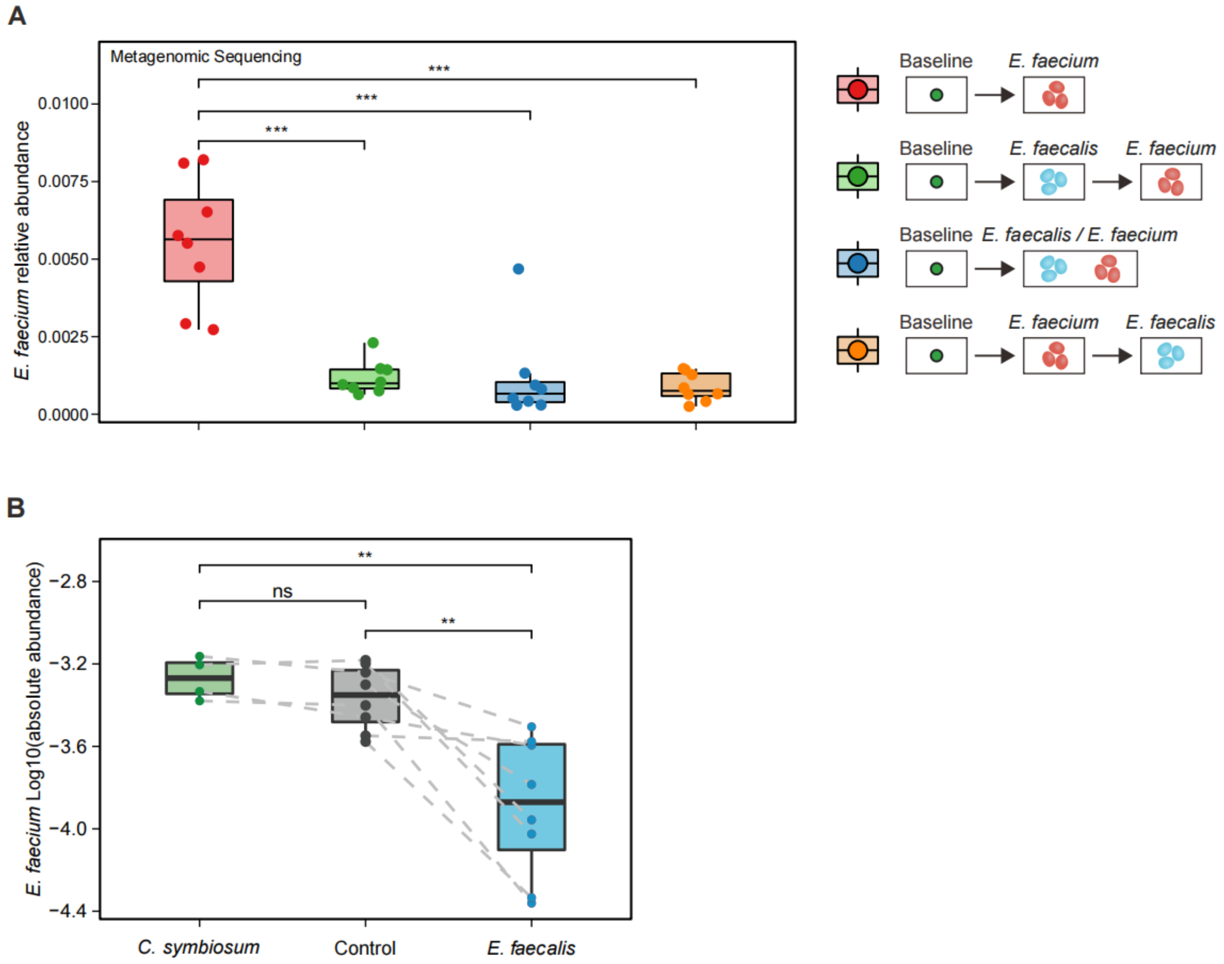


**Fig. S16.** ***E. faecalis*** **inhibits the growth of** ***E. faecium*** **in human stool-derived** ***in vitro*** **communities. (A)** The end-point abundance of *E. faecium*, measured by metagenomic sequencing. **(B)** The end-point abundance of *E. faecium* in communities inoculated with *E. faecalis* (inhibitory) or *C. symbiosum* (neutral) before *E. faecium* invasion (ns, not significant, ** $p < 0.01$, *** $p < 0.001$, Mann-Whitney U-tests).

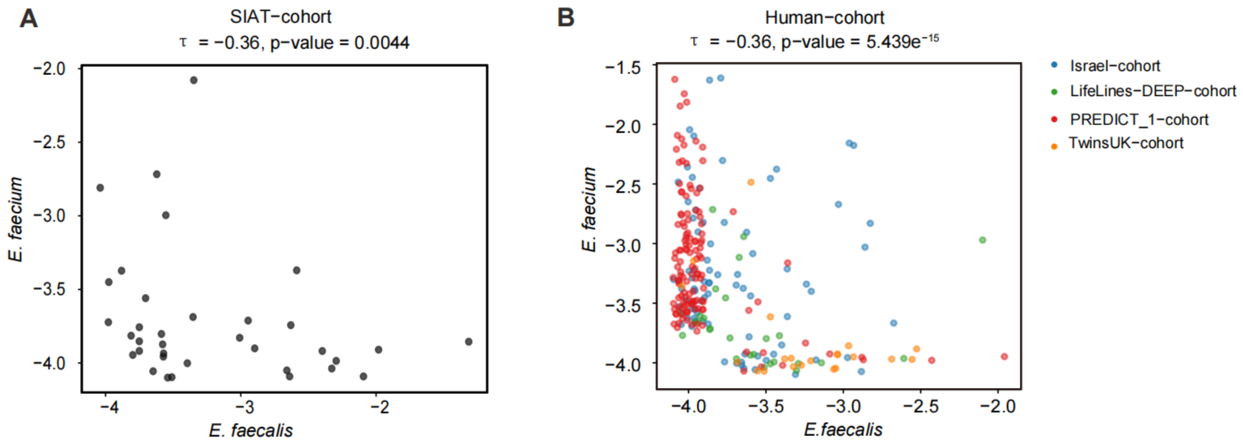


**Fig. S17. The relative abundance of *E. faecalis* and *E. faecium* is negatively correlated in human gut metagenomic samples. (A)** Negative correlation (Kendall correlation $\tau$ = -0.36) between the relative abundances of *E. faecalis* and *E. faecium* in the SIAT cohort. **(B)** Negative correlation (Kendall correlation $\tau$ = -0.36) between the relative abundances of *E. faecalis* and *E. faecium* in independent human cohorts. The detection limit in relative abundance was set to $10^{-4}$. 71.5% of the samples in the SIAT cohort and 93.8% of the samples in the four independent cohorts were negative (i.e., below the detection limit) for both *E. faecium* and *E. faecalis.*

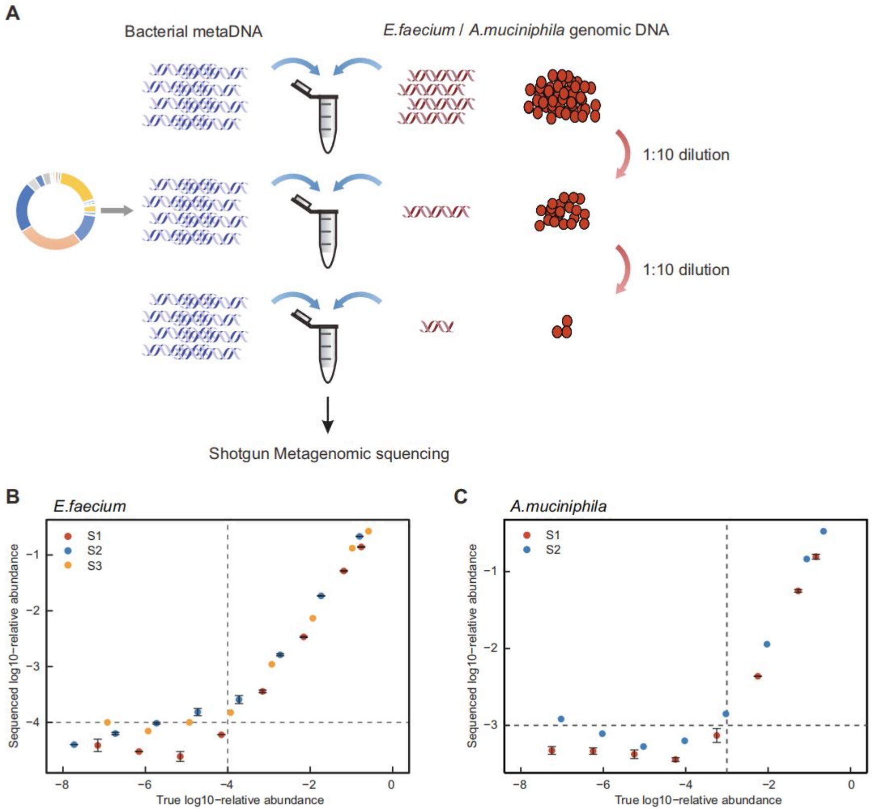


**Fig. S18. Quantification of the relative abundance of *E.faecium* and *A.muciniphila* by metagenomic sequencing. (A)** To confirm the accuracy of shallow metagenomic sequencing in quantifying the relative abundance of *E. faecium* and *A. muciniphila*, a spike-in experiment was conducted. The spike-in DNA of the target species (*E. faecium* or *A. muciniphila*) was 1:10 diluted for eight times and was added to the microbial metaDNA to a mixed DNA sample. The mixed DNA was then used for library construction and metagenomic sequencing. **(B-C)** By comparing the detected relative abundance generated by shallow metagenomic sequencing with the expected abundance, the accuracy and sensitivity of our workflow were determined. The detection threshold of *E. faecium* is 0.0001 and the detection threshold of *A. muciniphila* is 0.001.

**Table S1 Information of antibiotics used in this study.**

| Antibiotics | Name | Concentration（ug/ml） | Target | Class |
|---|---|---|---|---|
| Abx1 | Meropenem | 35 | Cell wall | beta-lactams |
| Abx2 | Cefoxitin | 10 | Cell wall | beta-lactams |
| Abx3 | Cefotaxime sodium salt | 40 | Cell wall | beta-lactams |
| Abx4 | Piperacillin Sodium+Tazobactam acid | 20+2.5 | Cell wall | beta-lactams |
| Abx5 | Vancomycin | 16 | Cell wall | Glycopeptides and lipoglycopeptides |
| Abx6 | Colistin sulfate salt | 20 | Cell wall | Miscellaneous agents |
| Abx7 | Ciprofloxacin | 12 | DNA synthesis | (Fluoro-)quinolones |
| Abx8 | Tobramycin | 400 | Protein synthesis | Aminoglycosides |
| Abx9 | Amikacin | 200 | Protein synthesis | Aminoglycosides |
| Abx10 | Erythromycin | 80 | Protein synthesis | Macrolides, lincosamides and streptogramins |
| Abx11 | Linezolid | 70 | Protein synthesis | Oxazolidinone |
| Abx12 | Tigecycline | 0.256 | Protein synthesis | Tetracyclines |

### References


1. An Illustrated Guide to Theoretical Ecology. *Journal of Mammalogy* **82**, 247-248 (2001).
2. P. Erdős, Alfréd Rényi, On the evolution of random graphs. *Publ. Math. Inst. Hung. Acad. Sci* **5**, 43 (1960).
3. S. Michel-Mata, X. W. Wang, Y. Y. Liu, M. T. Angulo, Predicting microbiome compositions from species assemblages through deep learning. i*Meta* **1**, (2022).
4. A. a. G. Paszke, Sam and Massa, Francisco and Lerer, Adam and Bradbury, James and Chanan, Gregory and Killeen, Trevor and Lin, Zeming and Gimelshein, Natalia and Antiga, Luca and Desmaison, Alban and Kopf, Andreas and Yang, Edward and DeVito, Zachary and Raison, Martin and Tejani, Alykhan and Chilamkurthy, Sasank and Steiner, Benoit and Fang, Lu and Bai, Junjie and Chintala, Soumith, PyTorch: An Imperative Style, High-Performance Deep Learning Library. *33rd Conference on Neural Information Processing Systems (NeurIPS 2019)*, (2019).
5. L. G. Buitinck L, Blondel M, Pedregosa F, Mueller A, Grisel O, Niculae V, Prettenhofer P, Gramfort A, Grobler J, Layton R, API design for machine learning software: experiences from the scikit-learn project. *arXiv preprint* **arXiv:1309.0238**, (2013).
6. L. Li *et al.*, An in vitro model maintaining taxon-specific functional activities of the gut microbiome. *Nature communications* **10**, 4146 (2019).
7. B. Javdan *et al.*, Personalized Mapping of Drug Metabolism by the Human Gut Microbiome. *Cell*, (2020).
8. H. P. Browne *et al.*, Culturing of 'unculturable' human microbiota reveals novel taxa and extensive sporulation. *Nature* **533**, 543-+ (2016).
9. B. Hillmann *et al.*, Evaluating the Information Content of Shallow Shotgun Metagenomics. *mSystems* **3**, (2018).
10. L. Maier *et al.*, Unravelling the collateral damage of antibiotics on gut bacteria. *Nature* **599**, 120-124 (2021).
11. T. EUCAST, European Committee on Antimicrobial Susceptibility Testing, Breakpoint tables for interpretation of MICs and zone diameters. *Version 5.0, 2015* (2015).
12. S. Huang *et al.*, Candidate probiotic Lactiplantibacillus plantarum HNU082 rapidly and convergently evolves within human, mice, and zebrafish gut but differentially influences the resident microbiome. *Microbiome* **9**, 151 (2021).
13. Z. Zhang *et al.*, Lactobacillus fermentum HNU312 alleviated oxidative damage and behavioural abnormalities during brain development in early life induced by chronic lead exposure. *Ecotoxicol Environ Saf* **251**, 114543 (2023).
14. P. D. Cani, W. M. de Vos, Next-Generation Beneficial Microbes: The Case of Akkermansia muciniphila. *Front Microbiol* **8**, 1765 (2017).
15. W. Xu *et al.*, Characterization of Shallow Whole-Metagenome Shotgun Sequencing as a High-Accuracy and Low-Cost Method by Complicated Mock Microbiomes. *Front Microbiol* **12**, 678319 (2021).
16. S. Dehbashi, H. Tahmasebi, P. Sedighi, F. Davarian, M. R. Arabestani, Development of high-resolution melting curve analysis in rapid detection of vanA gene, Enterococcus faecalis, and Enterococcus faecium from clinical isolates. *Trop Med Health* **48**, 8 (2020).
17. D. Zeevi *et al.*, Personalized Nutrition by Prediction of Glycemic Responses. *Cell* **163**, 1079-1094 (2015).
18. M. J. Bonder *et al.*, The effect of host genetics on the gut microbiome. *Nature Genetics* **48**, 1407-1412 (2016).
19. F. Asnicar *et al.*, Microbiome connections with host metabolism and habitual diet from 1,098 deeply

phenotyped individuals. *Nat Med* **27**, 321-332 (2021).

20. H. Xie *et al.*, Shotgun Metagenomics of 250 Adult Twins Reveals Genetic and Environmental Impacts on the Gut Microbiome. *Cell systems* **3**, 572-584 e573 (2016).
21. K. R. Clarke, Non‐parametric multivariate analyses of changes in community structure. *Australian Journal of Ecology* **18**, 26 (1993).
22. P. Dixon, VEGAN, a package of R functions for community ecology. *Journal of Vegetation Science* **14.6**, 3 (2003).
23. N. Segata *et al.*, Metagenomic biomarker discovery and explanation. *Genome Biology* **12**, R60 (2011).
24. T. E. Gibson, A. Bashan, H. T. Cao, S. T. Weiss, Y. Y. Liu, On the Origins and Control of Community Types in the Human Microbiome. *PLoS Comput Biol* **12**, e1004688 (2016).
